# Multifaceted effects of variable biotic interactions on population stability in complex interaction webs

**DOI:** 10.1101/2021.12.07.471695

**Authors:** Koya Hashimoto, Daisuke Hayasaka, Yuji Eguchi, Yugo Seko, Ji Cai, Kenta Suzuki, Koichi Goka, Taku Kadoya

## Abstract

Recent studies have revealed that biotic interactions in ecological communities vary over time, possibly mediating community responses to anthropogenic disturbances. This study investigated the heterogeneity of such variability within a real community and its impact on population stability in the face of pesticide application, particularly focusing on density-dependence of the interaction effect. Using outdoor mesocosms with a freshwater community, we found considerable heterogeneity in density-dependent interaction variability among links in the same community. This variability mediated the stability of recipient populations, with negative density-dependent interaction variability stabilizing whereas positive density-dependence and density-independent interaction variability destabilizing populations. Unexpectedly, the mean interaction strength, which is typically considered crucial for stability, had no significant effect, suggesting that how organisms interact on average is insufficient to predict the ecological impacts of pesticides. Our findings emphasize the multifaceted role of interaction variability in predicting the ecological consequences of anthropogenic disturbances such as pesticide application.

## Introduction

Predicting the magnitude to which anthropogenic disturbances affect ecosystems is essential for successful conservation and ecosystem management. In real ecosystems, all members of a biological community are linked to each other by biotic interactions such as predation, competition, and mutualism, and these interactions often cause unpredictable responses of biological communities to natural and/or anthropogenic disturbances (Rohr *et al*. 2006; Suttle *et al*. 2007; Doak *et al*. 2008). Indeed, ecologists have considered that community responses to disturbances highly depend on the strength of biotic interactions: weak interactions are likely to promote various types of community stability against system alterations (McCann *et al*. 1998; Wootton & Emmerson 2005; O’Gorman & Emmerson 2009; Kadoya *et al*. 2018). Notably, many earlier studies have postulated a ‘static view’ of interaction networks, focusing on long-term average properties of interactions responsible for community stability (May 1972; Dunne 2006; Allesina & Tang 2012). More recently, however, researchers have recognized that the effect of interactions can be highly variable due to behavioural or physiological trait plasticity and rapid evolution in response to intrinsic and extrinsic factors (Tylianakis *et al*. 2008; Ohgushi *et al*. 2012; Fussmann *et al*. 2014; McMeans *et al*. 2015; Ushio *et al*. 2018; Bartley *et al*. 2019). This view of variable interactions urges us to reconsider how biotic interactions determine community responses to anthropogenic disturbances.

One of the important sources of interaction variability is density-dependence in the *per capita* interaction effect. The *per capita* interaction effect is measured by changes in the *per capita* population growth of an interaction recipient species caused by slight changes in donor species density (Travis & Post 1979; Novak *et al*. 2016). The density-dependence of *per capita* interaction effects generates nonlinear functional responses that are well known to play a critical role in determining the stability of resource–consumer populations (Oaten & Murdoch 1975; Uszko *et al*. 2015), in which the variability in the *per capita* interaction effect is an implicit but essential underlying mechanism. Specifically, with negative interaction density-dependence, a decrease in the density of the recipient causes an interaction change that positively affects the recipient such that the collapse of recipient populations is avoided (i.e., stabilizing). Conversely, with positive interaction density-dependence, a decline in a recipient’s population provides a change in interaction effect disadvantageous to the recipient population, leading to a greater probability of extinction (i.e., destabilizing). Although the effects of functional response on population stability have been studied mostly in terms of resource–consumer relationships, the underlying mechanisms explicitly considering the variability in the interactions described above can be applied to other types of interactions, such as competition and mutualism (Holland *et al*. 2006; Kawatsu & Kondoh 2018).

An important challenge here is understanding the effects of interaction density-dependence in the context of large communities involving numerous species and a complex network of interactions among them. Recent theoretical studies have suggested that it is possible to scale up the stabilizing/destabilizing effects of interaction density-dependence from small numbers of species populations to communities consisting of many species (Kondoh 2003; Kawatsu & Kondoh 2018). At the same time, they have showed that incorporating heterogeneity of interaction density-dependence among links dramatically alters community stability and species diversity–stability relationships (Kawatsu & Kondoh 2018). In particular, the proportion and/or the effect size of stabilizing and destabilizing density-dependence are critical for determining the stability of the whole community because the stabilizing effects of species interactions may cancel out destabilizing interactions when they have a greater proportion and/or larger effects. Thus, quantifying heterogeneity in the forms of density-dependence among links is essential for understanding the stabilizing mechanisms of natural communities. However, describing the heterogeneity of the functional forms in natural communities is logistically challenging: this is because interaction density-dependence results from various mechanisms, such as constraints by prey handling time, optimal foraging, and adaptive defence and because these underlying mechanisms are shaped by the unique ecological and evolutionary context that each interaction link has experienced.

Here, we explored (i) heterogeneity in interaction density-dependence among interaction links within a real community and (ii) whether interaction density-dependence works to stabilize/destabilize populations in response to anthropogenic disturbances. We combined a manipulative open mesocosm experiment and nonlinear time-series analysis to track variability in interaction effects. As a model system, we chose pesticide and freshwater communities in paddy ponds, which are representative agricultural habitats in East Asia. We conducted three-year outdoor mesocosm experiments in which the ponds were fully crossed with two replicates each of an insecticide (fipronil) and an herbicide (pentoxazone). During the experimental period (approximately 140 days) of each year, we monitored the densities of ten community members, including phyto- and zooplankton, plants, and macroinvertebrates (Table S1), every two weeks. Based on the fortnight-interval data, we quantified interaction effects among these community members and their variability by empirical dynamic modelling (EDM), which is a recently developed analytical framework for nonlinear time-series data (Chang *et al*. 2017; Munch *et al*. 2020).

Using the EDM approach, we reconstructed the interaction network and its density-dependence of the communities in the paddies and distinguished observed interaction variability into three types in terms of recipient density-dependence in the *per capita* interaction effect (hereafter referred to as interaction density-dependence; IDD): (1) negative IDD, (2) positive IDD, and (3) density-independent interaction variability (see Methods for details). We then tested the following three specific predictions: (1) a negative IDD increases the stability of the recipient population because a decrease or increase in the recipient density is compensated for by changes in the interaction effect that are positive or negative to the recipient, respectively (Fig. 1a); (2) a positive IDD increases the instability of the recipient population because a decrease or increase in the recipient density is magnified by changes in the interaction effect that are negative or positive to the recipient, respectively (Fig. 1b); and (3) density-independent interaction variability is more likely to destabilize the recipient population because larger density-independent interaction changes induce greater unpredictable variability in the recipient population (Fig. 1c).

**Fig. 1.**
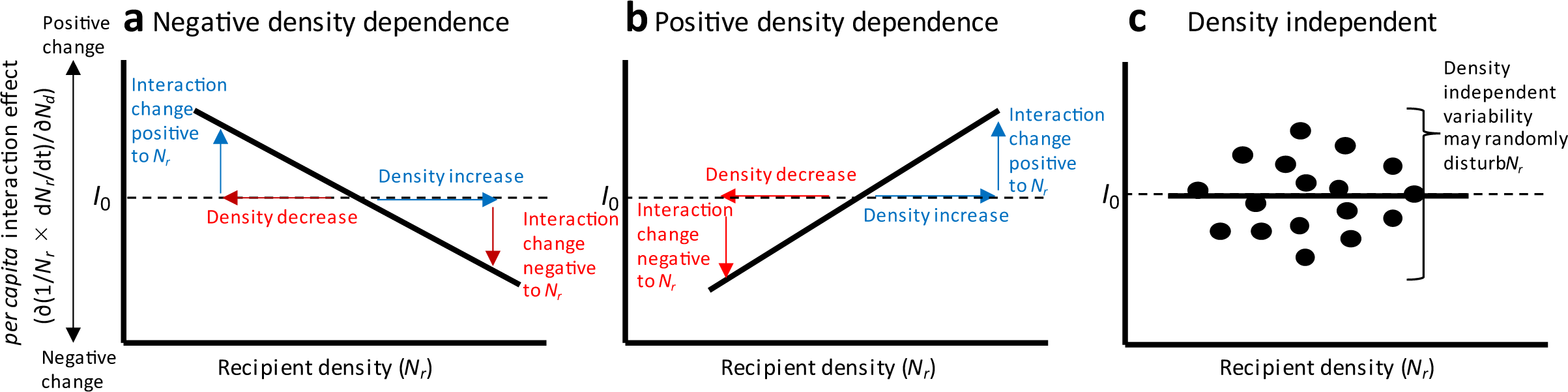
Schematic representation of three types of density-dependent (or density-independent) variability in the per capita interaction effect. *N_r_* and *N_d_* denote the recipient and donor densities, respectively. The *per capita* interaction effect (∂(1/*N_r_* × *dN_r_*/*dt*)/∂*N_d_*) may vary negatively or positively depending on recipient density or independent of recipient density. Red and blue arrows indicate density or interaction changes negative or positive to the recipient population, respectively. (a) Negative density-dependence. A decrease or increase in recipient density results in changes in the interaction effect that are positive or negative to the recipient, respectively, resulting in negative feedback and a greater likelihood of stabilizing the recipient population. (b) Positive density-dependence. A decrease or increase in density makes the interaction effect more negative or positive to recipient density, respectively, bringing positive feedback and potentially leading to population collapse or an outbreak. (c) Density-independent variability. Unlike density-dependent variability in interactions, density-independent interaction changes can cause positive and negative effects on recipient density in an unpredictable manner, resulting in a random disturbance to the recipient population. Note that the solid line in each panel intersects the *I*_0_ horizontal lines (*per capita* interaction effect = 0), i.e., the interaction effect changes from positive (negative) to negative (positive), but this is not necessary for interaction variability to have the potential to stabilizing or destabilizing effects. For example, stable coexistence between competitors may arise when competition, which is always negative, becomes stronger (in the downwards direction in the above figures) with increasing density.

## Results

### Pesticide impacts on the density of each community member

Here, we describe whether and how the three pesticide treatments (i.e., I: insecticide alone, H: herbicide alone, and I+H: insecticide and herbicide) affected the density of the ten community members in the experimental paddies. For the majority of the community members, insecticide applications had stronger impacts than herbicide applications on their density (Fig. 2a, Table S2). Insecticide treatment dramatically decreased the density of phytophilous and benthic predatory insects (i.e., Odonata larvae) [C (control) vs. I (insecticide alone)], which was consistent with previous studies (Hayasaka *et al*. 2012; Hashimoto *et al*. 2019). Additionally, although not statistically significant, the insecticide treatment decreased the density of detritivorous insects and increased the density of neustonic predatory insects and molluscs. The herbicide treatment significantly decreased only the density of macrophytes [C vs. H (herbicide alone)]. When we applied both the insecticide and the herbicide, phytoplankton, macrophytes, herbivores, phytophilous predators, and benthic predators significantly decreased in density, and molluscs significantly increased in density [C vs. I+H (mixture of insecticide and herbicide)]. The observed pesticide impacts may have been due to direct toxicity, indirect toxicity via biotic interactions, or a mixture of these factors.

**Fig. 2.**
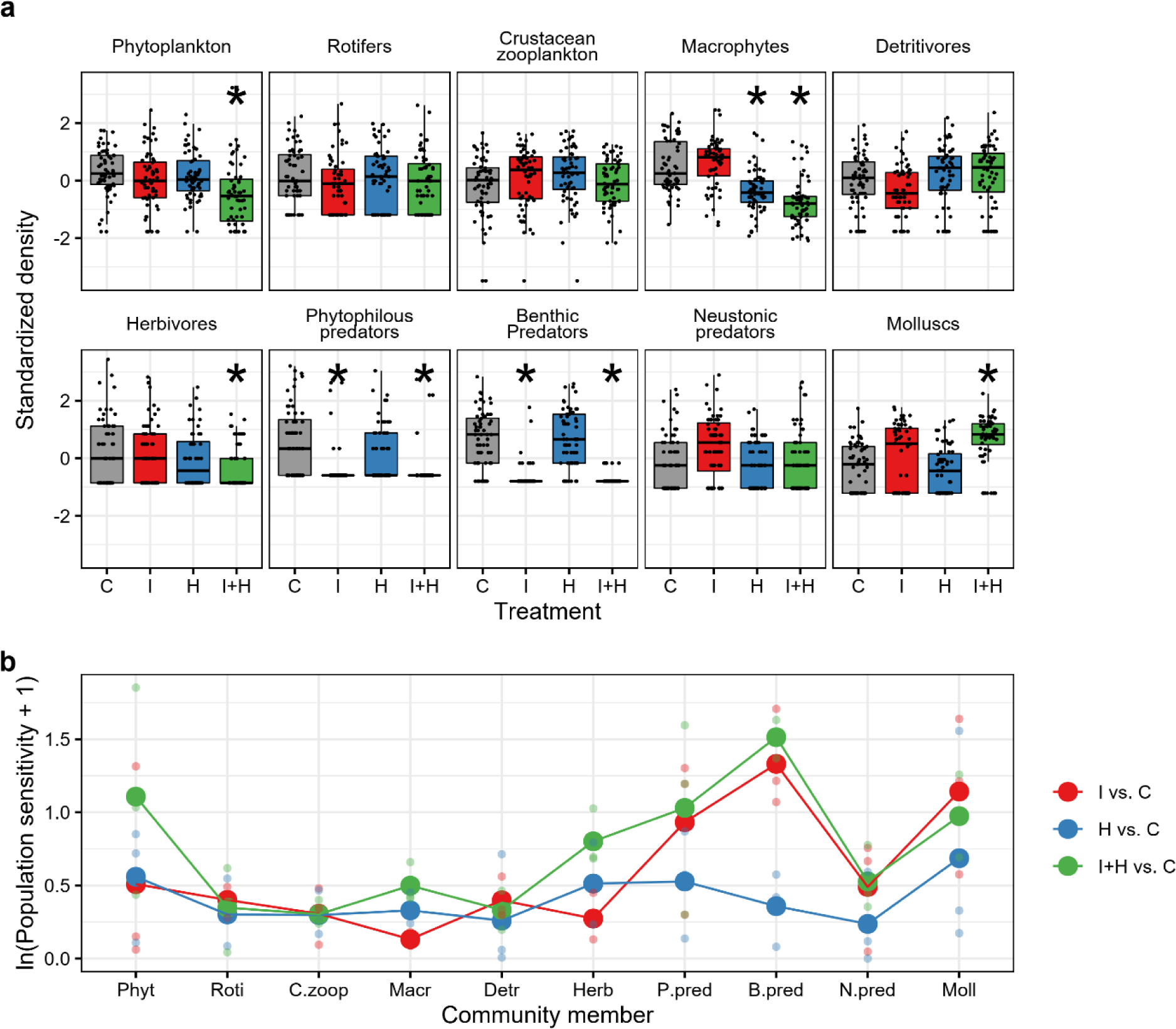
Pesticide impacts on the density of paddy community members. (a) Treatment effects on the density of each paddy community member. The treatment abbreviations are as follows: C: control, I: insecticide alone, H: herbicide alone, and I+H: mixture of insecticide and herbicide. Asterisks indicate significant differences compared to the control (α = 0.05). The midline, box limits, and whiskers indicate the median, upper and lower quartiles and 1.5× interquartile range, respectively. Points indicate the raw values. (b) Population sensitivity, which is measured as the absolute value of the log response ratio of the mean raw density in the pesticide treatments (either I, H or I+H) relative to that in the controls for each of the three experimental years. Large points indicate mean values, while small points indicate raw values. The abbreviations of the community members are as follows: Phyt: phytoplankton, Roti: rotifers, C.zoop: crustacean zooplankton, Macr: macrophytes, Detr: detritivores, Herb: herbivores, P.pred: phytophilous predators, B.pred: benthic predators, N.pred: neustonic predators, and Moll: molluscs.

We evaluated population sensitivity to pesticide treatments, which is a proxy of population instability, by calculating the absolute value of the log response ratio (LRR) of the mean density in a pesticide treatment relative to that in the controls (Hedges *et al*. 1999). We calculated LRR*_i_*_,*j*_ for community member *i* in year *j* for every pesticide treatment as ln((*T_i_*_,*j*_ + 0.1)/(*C_i_*_,*j*_ + 0.1)), where *T_i_*_,*j*_ is the mean density in either the I, H, or I+H treatment and *C_i_*_,*j*_ is the mean density of the controls. We added 0.1 to the numerator and denominator because there were several zero data points for both *T_i_*_,*j*_ and *C_i_*_,*j*_ (Martinson & Raupp 2013). We found that there was considerable variation in sensitivity among community members (Fig. 2b). That is, some members were sensitive (unstable), and other members were relatively less sensitive (stable) to pesticide disturbance. In particular, rotifers, crustacean zooplankton, and detritivores were relatively less sensitive to all three pesticide treatments. Additionally, this variation in the sensitivity among different community members changed depending on the three pesticide treatments. Phytophilous, benthic, and neustonic predators and molluscs were more sensitive to the I and I+H treatments than to the H treatment, while macrophytes and herbivores were more sensitive to the H and I+H treatments than to the I treatment.

### Heterogeneity in interaction variability among different links

Our EDM analyses successfully reconstructed the interaction network among paddy community members in each treatment (Fig. 3, Table S3, Fig. S2). Fig. 3a shows the interaction networks in the controls. Note that the arrows represent the net effects of interactions (i.e., the sum of direct and indirect effects) rather than the sole direct effects. Most of the interactions were biologically interpretable; for instance, phytoplankton positively affected rotifers, and rotifers negatively affected phytoplankton but positively affected crustacean zooplankton, suggesting prey–predator interactions (Fig. 3a, APPENDIX 3). As several previous studies have indicated (Wootton & Emmerson 2005), the distribution of the mean interaction strength given no disturbances was skewed towards a weaker strength (Fig. 3b), and this tendency did not change for the pesticide application treatments (Fig. S2). For every treatment, the numbers of negative and positive interactions were similar (Fig. S2). Furthermore, the interaction effect varied over time within each year even without pesticides (Fig. 3c, Fig. S3). Notably, the magnitude of interaction temporal variability differed greatly among interaction pairs (Fig. 3c). For example, the interaction effect on herbivorous insects showed only subtle variability, but that on molluscs greatly varied over time (Fig. 3c).

**Fig. 3.**
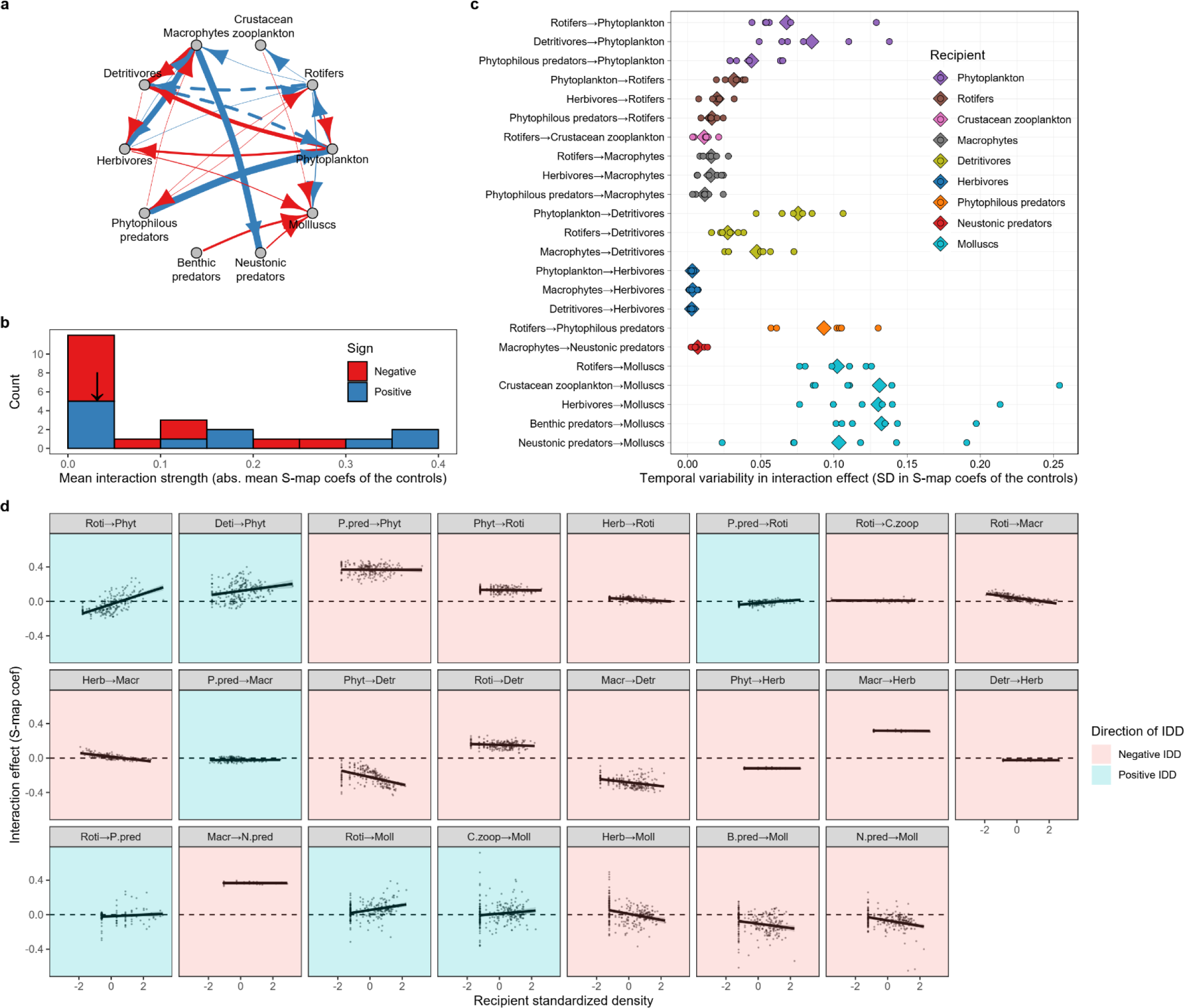
Heterogeneous variability in *per capita* interaction effects within the experimental paddy community. (a) Interaction networks of the controls (i.e., treatments without pesticide disturbances) reconstructed by EDM analysis. Red and blue arrows indicate negative and positive interactions, respectively. Their thickness is proportional to the mean interaction strength represented by the absolute values of the mean S-map coefficient averaged over all the experimental periods and replicates. Solid arrows: *P* < 0.05, dashed arrows: 0.05 < *P* < 0.1. (b) Right-skewed distribution of the mean interaction strength in the control treatments (absolute values of mean S-map coefficients averaged over all the experimental periods and replicates in the controls). The vertical arrow indicates the median values. (c) Temporal variability in the *per capita* interaction effect measured by the SDs of the S-map coefficients over the experimental periods in the control treatment. Diamonds and circles indicate the overall means and raw data per replicate per year, respectively. (d) Density-dependence in the *per capita* interaction effect (IDD). For each interaction pair, S-map coefficients at each time point (grey dots) were plotted against the recipient standardized density at the corresponding time point. Solid segments represent simple linear regression lines with 95% confidence intervals (grey shading). The direction of the regression slopes (i.e., negative or positive interaction density-dependence (IDD)) is indicated by background colours: red, negative IDD; and blue, positive IDD. The abbreviations of the community members are as follows: Phyt: phytoplankton, Roti: rotifers, C.zoop: crustacean zooplankton, Macr: macrophytes, Detr: detritivores, Herb: herbivores, P.pred: phytophilous predators, B.pred: benthic predators, N.pred: neustonic predators, and Moll: molluscs.

By plotting the interaction effect against the recipient density at each time point, we found that there was considerable heterogeneity in the IDD (i.e., recipient density-dependence in interaction effect) among different interaction links in terms of not only its steepness but also its direction. The number of negative slopes (potentially stabilizing) was twice the number of positive slopes (potentially destabilizing) (Fig 3d). Furthermore, density-independent interaction variability, represented by the deviation of the raw interaction effect data from the regression slopes in Fig. 3d, also differed considerably among different interaction links even when compared between similar slope steepnesses.

### Effects of interaction density-dependence on population sensitivity

We examined the effects of three types of interaction density (in)dependence on recipient population sensitivity to the three pesticide treatments by using a multiple regression approach under the controls of mean interaction strength and recipient functions (Table 1, Fig. 4). Note that the effects of temporal variability shown in Table 1 and Fig. 4b, f, j were under the control of the dependence of interaction on recipient density, thereby representing the effects of the recipient-density-*independent* interaction variability. We observed that negative IDD tended to have negative effects on recipient sensitivity to pesticide disturbance (i.e., stabilizing effects) (Fig. 4a, e, i), whereas positive IDD was more likely to have neutral or positive effects on sensitivity (i.e., destabilizing effects) (Fig. 4a, e, i). The likelihood ratio *χ*^2^ of the interaction term between IDD × direction (negative or positive) of IDD was relatively high, although it was not statistically significant for the I and H treatments (Table 1). For every pesticide treatment, we observed positive effects of interaction temporal variability on recipient sensitivity to all pesticide treatments (Fig. 4b, f, j), suggesting that density-independent interaction variability was destabilizing. The interaction temporal variability had the greatest likelihood ratio *χ*^2^ (Table 1), suggesting that it was the most important variable for every treatment. Although weak interactions are considered stabilizing forces of communities, the mean interaction strength did not have a significant effect on recipient sensitivity to any pesticide treatment (Table 1, Fig. 4c, g, k).

**Fig. 4.**
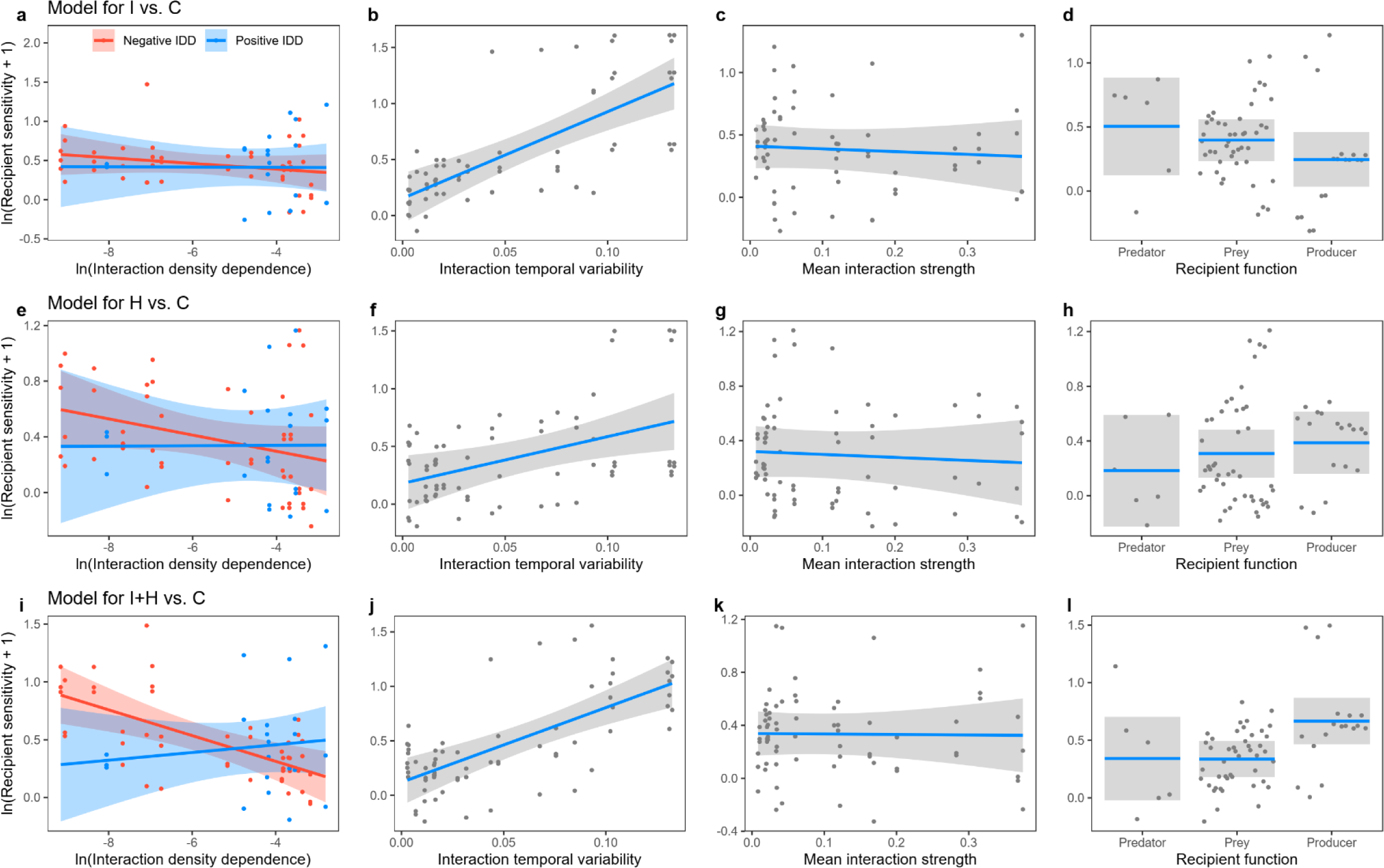
Stabilizing and destabilizing effects of different interaction properties on recipient population sensitivity to the three pesticide treatments. The insecticide treatment (I vs. C, a-d), herbicide treatment (H vs. C, e-h), and insecticide + herbicide treatment (I+H vs. C, i-l) are shown. Downwards (upwards) effects can be interpreted as stabilizing (destabilizing) for recipient density. The effects of interaction density-dependence (IDD) (a, e, i), interaction temporal variability (b, f, j), mean interaction strength (c, g, k), and recipient function (d, h, l) are shown. In each panel, fitted lines with 95% confidence intervals were obtained with all covariates held at median values. This indicates that the effects of the mean interaction strength and interaction temporal variability visualized in the figure are independent of density-dependent variability in interaction effects. In Panels a, e, and i, the direction of IDD is shown in red (negative IDD) and blue (positive IDD). The grey, red, and blue dots are partial residuals. A statistical summary of the models is shown in Table 1.

**Table 1.** Relative importance of different interaction properties on recipient population sensitivity to the insecticide treatment (a, I vs. C), the herbicide treatment (b, H vs. C), and the insecticide + herbicide treatment (c, I+H vs. C) Likelihood ratio tests based on LMMs were conducted. Bold indicates statistical significance.

| (a) I vs. C |  |  |  |
| --- | --- | --- | --- |
| | likelihood ratio $\chi^2$ | df | P |
| Mean interaction strength | 0.28 | 1 | 0.59 |
| Interaction temporal variability | **26.78** | **1** | **< 0.001** |
| Interaction density-dependence (IDD) | 0.35 | 1 | 0.55 |
| Direction of IDD (negative or positive) | 0.03 | 1 | 0.86 |
| IDD × direction of IDD | 0.41 | 1 | 0.52 |
| Recipient function | 2.88 | 2 | 0.24 |
| (b) H vs. C |  |  |  |
| | likelihood ratio $\chi^2$ | df | P |
| Mean interaction strength | 0.25 | 1 | 0.62 |
| Interaction temporal variability | **7.62** | **1** | **0.01** |
| Interaction density-dependence (IDD) | 0.68 | 1 | 0.41 |
| Direction of IDD (negative or positive) | 0.07 | 1 | 0.8 |
| IDD × direction of IDD | 1.06 | 1 | 0.3 |
| Recipient function | 1.23 | 2 | 0.54 |
| (c) I+H vs. C |  |  |  |
| | likelihood ratio $\chi^2$ | df | P |
| Mean interaction strength | 0.01 | 1 | 0.92 |
| Interaction temporal variability | **21.9** | **1** | **< 0.001** |
| Interaction density-dependence (IDD) | 1.61 | 1 | 0.21 |
| Direction of IDD (negative or positive) | 0.12 | 1 | 0.73 |
| IDD × direction of IDD | **7.5** | **1** | **0.01** |
| Recipient function | **9.96** | **2** | **0.01** |

## Discussion

Our study clearly demonstrated that different types of interaction density (in)dependence certainly coexisted in the same community in the experimental paddies and that the interaction variability mediated the stability of the corresponding recipient populations (i.e., populations receiving interaction effects) embedded in the complex interaction webs. Consistent with our expectations, the different types of interaction variability had contrasting effects on population stability; negative IDD tended to decrease population sensitivity to pesticides (stabilizing), whereas positive IDD was more likely to magnify it (destabilizing). Previous studies have provided mixed predictions regarding the effects of interaction variability on population and/or community stability in response to external disturbances; some studies suggested improving effects (Kondoh 2003; Navarrete & Berlow 2006; Rooney *et al*. 2006, 2008; Loeuille 2010), while others suggested impairing effects (Rooney *et al*. 2006; Calizza *et al*. 2017). In this context, we provided empirical evidence that whether interaction variability is stabilizing or destabilizing depends on the type of variability (i.e., how the *per capita* interaction effect responds to density changes), which was previously suggested only by theoretical works (Kawatsu & Kondoh 2018; Kawatsu 2020). Furthermore, we found that not only positive IDD but also density-independent interaction variability consistently increased population sensitivity to pesticide treatments (destabilizing). We argue that incorporating the effects of density-independent interaction variability, or stochasticity in interaction strength (Shoemaker *et al*. 2020), into the study of dynamic interaction networks is a promising avenue for future research.

The observed IDD was mostly biologically interpretable, especially in cases of seemingly prey–predator interactions. For example, the negative interaction effect of benthic predators (dragonfly larvae) and neustonic predators (water striders) on their potential prey molluscs became stronger with increasing density of molluscs (Fig. 3d; B.pred→Moll and N.pred→Moll), suggesting that these apex predators change their foraging behaviour in response to changes in prey density such that they choose more abundant prey. Such adaptive foraging in response to changes in prey density is regarded as a stabilizing force in predator‒prey interactions (Kondoh 2003; Rooney *et al*. 2006, 2008; Loeuille 2010; but see Abrams 2010). On the other hand, the negative effect of phytophilous predators (damselfly larvae) on their potential prey (rotifers) was weaker with increasing rotifer density (Fig. 3d; P.pred→Roti), which was potentially destabilizing. Such density-dependence can arise from the saturating predation rate due to the constraints of predator handling time (Holling 1959; Uszko *et al*. 2015). Unlike those interactions that were easily interpretable, there were interaction links whose density-dependence was hard to interpret. For example, the reasons why the negative interaction effect of phytoplankton on detritivores was negatively density dependent (Fig. 3d; Phyt→Detr) and why the positive interaction effect of rotifers on macrophytes was negatively density dependent (Fig. 3d; Roti→Macr) are unknown. Notably, the literature has shown that we still know little about functional forms of density-dependence in positive interaction effects such as the exploitation of benefits by a predator species from its prey (Norris & Johnstone 1998) and mutualistic effects (Holland *et al*. 2002). Our analysis sheds new light on never-detected IDD in real ecosystems, even though the specific underlying mechanisms are not yet clear. Identifying mechanisms may require further research on a theory that incorporates evolutionary processes in developing IDD (Kawatsu & Kondoh 2018).

Although there has been growing consensus that weak interactions play a significant role in community stability (McCann *et al*. 1998; Kadoya & McCann 2015; Kadoya *et al*. 2018), in our study, we did not find any stabilizing effects of mean interaction strength (Fig. 4c, g, k). However, it is possible that interaction strength can influence population stability via the associations between mean interaction strength and interaction variability (McCann 2000). Berlow (1999) observed negative relationships between mean interaction strength and spatial variability, in which some weak interactions on average were extremely variable among spatial replicates, but strong interactions were less variable. The author argued that the variability of weak interactions should play an important role in community and ecosystem organization. Our additional analysis revealed a similar pattern, an overall negative association between the mean interaction strength and interaction temporal variability (Fig. S4a), although the trends were not statistically significant, probably due to data variance heterogeneity (Fig. S4a). Interestingly, the pattern was mainly driven by negative density-dependent interactions, which were proven to be stabilizing (Fig. S4b). As such, we suggest that future studies exploring the relationships between interaction strength and variability, including both density-dependent and density-independent interactions, are required to draw a complete picture of how those processes act interactively to determine community stability.

### Methodological considerations

When interpreting our results, which were obtained from a combination of manipulative mesocosm experiments and EDM methods, some caution is needed. First, although there have been an increasing number of reports of EDM methods being applied in ecological studies (Kawatsu & Kishi 2018; Matsuzaki *et al*. 2018; Ushio *et al*. 2018; Rogers *et al*. 2020; Kawatsu *et al*. 2021), whether EDM can indeed uncover true causality from real data is still unclear (Cobey & Baskerville 2016; Barraquand *et al*. 2021) for several reasons, such as violation of the assumptions of EDM on the stationarity of time series (Munch *et al*. 2020), synchronization among multiple time series due to seasonality, which are known to inhibit causal inferences by EDM (Cobey & Baskerville 2016), and missing variables (but see Wang *et al*. 2020). In our study, we performed a seasonal surrogate analysis (see Methods) to minimize the confounding effect of seasonality, although we recognize that the results of this analysis should be interpreted carefully (Baskerville & Cobey 2017; Sugihara *et al*. 2017). Nevertheless, most of the interactions detected by CCM and quantified by S-map in our study were biologically reasonable (Fig. 3a, APPENDIX 3).

Second, estimating interaction effects by S-map models has been considered sensitive to the choice of variables (Ushio *et al*. 2018; Munch *et al*. 2022). This becomes more serious when the number of interacting variables is large because, in such a case, potential combinations of variables become increasingly larger, leading to unstable quantification of interaction effects (Chang *et al*. 2021). In our study, interaction effects were measured among the functional groups, not among the species. Although this simplification may be misleading because temporal species turnover within these groups could affect the observed interaction effects (Ives *et al*. 1999), it reduced the number of variables and hence the number of potential variable combinations. Thus, our estimation of interaction effects may be relatively robust. In addition, our observations were made over a two-week interval, and the interaction effects fluctuated over this relatively short time scale (Fig. S3b). If such a time scale was actually shorter than the rate of temporal species turnover within functional groups, the effects of temporal changes in species composition within functional groups on the observed interaction variability may have been relatively minor.

Finally, the number of replicates used in our experiment was low (i.e., 2). Previous ecotoxicological, mesocosm studies have also used a relatively small number of replicates (ranging from 2–5), probably due to resource and/or effort limitations (Hashimoto *et al*. 2019). Further studies are needed to confirm that the observed interaction dynamics are reproducible. Nevertheless, our additional analysis confirmed that the main source of variation (in a statistical sense) in the interaction effect was census date and treatment, not residuals (Fig. S5). Therefore, we believe that the dynamics of the interaction effect were relatively similar between the two replicates.

### Conclusion

The ecological impacts of pesticides on multispecies communities include direct toxicity and indirect toxicity mediated by biotic interactions (Relyea & Hoverman 2006; Rohr *et al*. 2006; Hashimoto *et al*. 2019). Our study emphasized the importance of the latter aspects of toxicity. Importantly, although the processes by which toxicity is mediated by interaction variability are apparently complex, some of the observed effects of the pesticides were consistent with previous studies. For example, the destructive effects of the insecticide fipronil on Odonata larvae (phytophilous and benthic predators in our study) and associated changes in community composition have been repeatedly reported in recent studies (Hayasaka *et al*. 2012; Kasai *et al*. 2016; Hashimoto *et al*. 2019; Ishiwaka *et al*. 2024). Community-level effects of the herbicide pentoxazone have not been well tested, but our study showed that negative effects of pentoxazone on macrophytes were consistently observed across the three experimental years. These results imply that even the causes and consequences of such established effects of these pesticides may involve density-(in)dependent interaction variability. This knowledge of the underlying mechanism is critical if we need to predict the impacts beyond the phenomenological understanding of them.

Recently, there has been a growing body of theoretical developments and new empirical studies on the species coexistence (Chesson 2000, 2018; Barabás *et al*. 2018; Yamamichi *et al*. 2022) and the stability of ecological communities without assuming equilibrium (Ushio *et al*. 2018; Cenci & Saavedra 2019). Understanding general laws about species diversity and stability in nonequilibrium communities would contribute to the understanding of the long-term effects of anthropogenic disturbances (Kondoh *et al*. 2020; Hallett *et al*. 2023; Ushio *et al*. 2023). We suggest several promising directions for future studies. First, although field studies on nonequilibrium communities are not easy, they can be achieved by recently developed techniques such as the EDM framework. By combining this new technique with knowledge about the roles of interaction variability in determining community stability, we obtained a tool set to predict which communities are most sensitive to anthropogenic disturbances. Our study provides the first step in using the EDM framework effectively for such purposes. Second, scaling up from population stability to whole-community stability is needed to generalize our results. Our study examined population-level stability, not community-level stability. There are theoretical studies that have explored the role of density-dependence in *per capita* interaction effects on whole-community stability (Kawatsu & Kondoh 2018; Kawatsu 2020), but these studies did not account for density-independent interaction variability, which was found to be consistently destabilizing in our study. Furthermore, to our knowledge, there are no empirical studies on this topic. Scaling up to whole-community stability with full consideration of density-(in)dependent interactions would be a promising task for future studies. Compiling such studies could serve to improve novel ecological risk assessments, contributing to effective ecosystem management in the face of ongoing environmental changes due to human activities.

## Materials and methods

### Experimental design

This study is a continuation of our previous study (Hashimoto *et al*. 2019), which examined the single and combined effects of an insecticide and an herbicide on freshwater communities in paddy mesocosms. We chose fipronil and pentoxazone as test chemicals because both are commonly used in Japanese crop management, and their physicochemical properties are as typical as the properties of pesticides used in Japanese paddies (Japan Plant Protection Association 2011). Fipronil acts upon the central nervous systems of animals by inhibiting GABA receptors, which are dominant in the nervous systems of arthropods (Gant *et al*. 1998). It is relatively stable in water and easily adsorbed to sediment, where it can persist for long periods (Simon-Delso *et al*. 2015). Pentoxazone is an inhibitor of chlorophyll biosynthesis (Hirai *et al*. 2001). It is quickly degraded in water by hydrolysis but is readily adsorbed to sediment, where it can persist for a relatively long time (Japan Plant Protection Association 2011; Nagai *et al*. 2011). Importantly, the direct toxicity of these chemicals is selective; fipronil is highly toxic to arthropods but moderately toxic to other animals and plants, while pentoxazone suppresses many vascular plants, but its toxicity to animals is low.

The experimental procedures used were described in Hashimoto *et al*. (2019). In March 2017, we buried eight independent fibre-reinforced plastic (FRP) tanks (280 cm length × 120 cm width × 40 cm depth, RK 3014, KAISUIMAREN Co., Ltd., Toyama Prefecture, Japan) in the ground on the campus facilities of Kindai University (Nara Prefecture, Japan). Then, we spread sediments from uncontaminated areas near the study site on the bottom of each tank (to a depth of approximately 30 cm). Each tank was randomly assigned to one of four treatments, i.e., C: controls, I: insecticide alone, H: herbicide alone, and I+H: mixture of insecticide and herbicide.

Pesticide applications were performed at the beginning of the growing season of each experimental year (2017–2019), i.e., once a year. Specifically, we repeated the following procedure each year. In mid-April, the mesocosms were flooded with dechlorinated water to a depth of approximately 5 cm. From the end of May to early June, we transplanted insecticide-treated and control rice seedlings (Hino-hikari variety) via an array of 4 × 10 at 25 cm intervals. We applied an insecticide (fipronil) and an herbicide (pentoxazone) in the same way as recommended for commercial rice fields. We treated nursery boxes of rice seedlings with Prince® (1% granular fipronil, HOKKO Chemical Industry, Inc., Tokyo, Japan) at a rate of 50 g/box 24 h prior to transplanting. Immediately after transplanting the rice seedlings, we applied Sainyoshi Flowable^®^ (8.6% pentoxazone, KAKEN Pharmaceutical Co., Ltd., Tokyo, Japan) at a rate of 1.7 mL/tank (i.e., 500 mL/10-a) to the mesocosms. We confirmed that these pesticide applications succeeded by monitoring the temporal dynamics of the water and sediment concentrations of the insecticide and herbicide (Fig. S1). The experiment was terminated in mid-October (i.e., approximately 140 days). We monitored the densities of ten functional groups in the community (eukaryotic phytoplankton, rotifers, crustacean zooplankton, macrophytes, detritivorous insects, herbivorous insects, phytophilous predatory insects, benthic predatory insects, neustonic predatory insects and molluscs; see Table S1) every two weeks throughout the approximately 140-day experimental period in each experimental year. Aggregating species by functional groups has typically been adopted as a strategy for describing aquatic food webs (Ives *et al*. 1999, 2003; Benincá *et al*. 2008; Matsuzaki *et al*. 2018). The detailed monitoring methods are described in APPENDIX 1. As such, we acquired 8 tanks × 3 years = 24 fragments of time series, each of which had 10 consecutive timepoints, for each of the 10 community members.

### Data processing for EDM analysis

All the statistical analyses were performed using the statistical environment ‘R’ Version 3.6.3 (R Core Team 2020). Before all the analyses, the time series of density data were normalized by ln(*x* + 1) transformation (except for macrophytes) and then standardized to zero means and unit variances to facilitate comparisons among different taxa. Standardization was performed for all 24 fragments of time series data as a composite. Such data processing has been recommended in several EDM protocols (Sugihara *et al*. 2012; Deyle *et al*. 2016; Chang *et al*. 2017).

### Pesticide impacts on community member density

We examined the effects of pesticide application on the density of each member by using linear mixed models (LMMs) with the package ‘glmmTMB’ Version 1.0.2.1 (Brooks *et al*. 2017). LMMs included standardized density as a response variable and treatment and census week (categorical) as explanatory variables. The random parts of the models were as follows: we specified tank identity and year as random intercepts and assumed an AR-1 temporal correlation structure within a given tank within a year. The statistical significance of the fixed terms was tested by Type III likelihood ratio tests, followed by Dunnett-type post hoc pairwise comparisons (two-sided) with the package ‘emmeans’ Version 1.5.2.1 (Lenth 2020).

### Calculation of population sensitivity to pesticides

To evaluate population sensitivity to the three pesticide treatments (as a proxy of population stability in response to pesticide disturbances), we used absolute values of the log response ratio (LRR). The LRR quantifies the proportional effects of experimental treatments on population density on a natural logarithmic scale, with positive values indicating greater population density in the treatment tanks and negative values indicating the opposite. We calculated LRR*_i_*_,*j*_ for community member *i* in year *j* for every pesticide treatment as ln((*T_i_*_,*j*_ + 0.1)/(*C_i_*_,*j*_ + 0.1)), where *T_i_*_,*j*_ is the mean density in either I, H, or I+H treatment and *C_i_*_,*j*_ is the mean density of the controls. We added 0.1 to the numerator and denominator because there were several zero data points for both *T_i_*_,*j*_ and *C_i_*_,*j*_ (Martinson & Raupp 2013). We used the absolute values of the LRR as the population sensitivity because our objective was to evaluate the stability of the populations rather than to determine whether the population increased or decreased in response to the treatments.

### Empirical dynamic modelling

To evaluate the effects of biotic interactions within the communities in the experimental paddies, we performed EDM analyses. The essence of EDM is a reconstruction of the true dynamics of a system by a subset of the variables of that system, which is called state-space reconstruction or attractor reconstruction. EDM was originally designed to analyse single, relatively long time series (e.g., at least > 35–40 consecutive time points; Sugihara *et al*. 2012), yet several alternative approaches have been suggested for analysing shorter time series by combining multiple time series that are too short to be analysed individually (Hsieh *et al*. 2008; Clark *et al*. 2015). Following these approaches, we assembled time series from the different experimental paddies and different years, which consisted of 10 time points × 8 paddies × 3 years = 240 time points in total, and analysed these assembled time series as a whole for each of our EDM analyses. Specifically, we allowed a single reconstructed attractor to consist of the whole data of every replicate, treatment and year, while we constrained every single data point on the attractor to be generated only from the data of the same replicate in the same year. This approach implicitly assumes that the time series of different replicates, treatments and years share a common dynamic rule.

We first detected dynamic causality by performing convergent cross-mapping (CCM) (Sugihara *et al*. 2012) to identify interacting pairs of community members and determine their interaction directions. Then, we constructed a multivariate S-map (Deyle *et al*. 2016) to track the time-varying, *per capita* interaction effect among interacting pairs determined by CCM. The detailed methods are described in APPENDIX 2. All EDM analyses were performed by using the package ‘rEDM’ Version 0.7.5 (Ye *et al*. 2020).

#### Detection of causalities by CCM

CCM is a recently developed analysis used to test causalities between two variables that potentially interact with each other. In short, CCM examines the correspondence among reconstructed attractor manifolds to test causalities among variables. We tested the causalities of all pairs of 10 community members (Tables S1, S3).

#### Tracking interaction variability via multivariate S-map

To estimate the *per capita* interaction effect among community members in the experimental paddies at each time point, we performed a multivariate S-map. The multivariate S-map is a locally weighted sequential linear regression at each location of an attractor manifold reconstructed by multivariate embedding. Deyle *et al*. (2016) proposed that regression coefficients of the S-map (‘S-map coefficients’) at each time point can be interpreted as time-varying interaction effects. For an S-map model to predict community member A, the estimated S-map coefficients of community member B are regarded as the time-varying interaction effect of community member B on A. To determine the interaction effect at the *per capita* level, we took the *per capita* population growth rate as the response variable (Suzuki *et al*. 2017). The S-map coefficients estimated in this way theoretically correspond to the first-order partial derivatives of a recipient’s *per capita* growth rate on a donor density: ∂(1/*N_r_* × *dN_r_*/*dt*)/∂*N_d_*, where *N_r_* and *N_d_* denote the density of the focal recipient and donor, respectively (Travis & Post 1979; Novak *et al*. 2016). The *per capita* growth rate was approximated by calculating ln(*N_t_*_+1_/*N_t_*), where *N_t_* and *N_t_*_+1_ are the raw density data of the focal species at time points *t* and *t*+1, respectively. Since there were several zero values in the raw density data, one was added to every density data point to calculate the *per capita* growth rate. Note that S-map is sensitive to data with stochastic noise, and we used a modified version of S-map, namely, regularized S-map, which incorporates a penalty term to avoid overfitting (Cenci *et al*. 2019). Regularized S-map has been shown to provide relatively robust estimations of interaction effects (Cenci *et al*. 2019). As S-map analyses were performed for reconstructed manifolds embedded by using the whole data of eight paddies and three years as a composite (i.e., 240 time points), we obtained interaction effects not only at every time point but also for every paddy (and thus every treatment) and every year (Fig. S3). To confirm whether the dynamics of the interaction effect were consistent among replicates, we performed two-way ANOVA for every interaction link, examining the effects of treatment, census date and their interaction on S-map coefficients. For simplicity, we did not account for any random effects or temporal autocorrelation.

### Effects of interaction density-dependence on population sensitivity

We performed multiple regressions to determine the effects of the interaction density-dependence (IDD) of each interaction link on the sensitivity of the corresponding recipient population. For this purpose, we first calculated the three interaction properties, IDD, interaction temporal variability and mean interaction strength.

IDD was represented by the absolute values of regression slopes between S-map coefficients and the standardized density of recipient populations. To do this, simple linear regressions using all treatment data over the three-year experimental period were performed for each interaction link (see Fig. 3d), and the absolute values of the regression coefficients for all the interaction links were obtained. Regardless of whether the direction of the slopes was negative or positive, higher absolute values of regression coefficients indicate stronger density-dependence in interaction effects (i.e., more variable interactions). The mean interaction strength or interaction temporal variability was calculated by first computing the mean or standard deviation of the S-map coefficients of the controls over the experimental period per year per replicate and then averaging their absolute values over all the years and replicates.

We constructed linear mixed models (LMMs) with the package ‘glmmTMB’ Version 1.0.2.1 (Brooks *et al*. 2017). The LMMs included ln(*x* + 1)-transformed absolute values of LRR (population sensitivity) as a response variable and all three interaction properties as explanatory variables. The IDD was ln(*x*)-transformed to improve the model fit. In addition, to test whether the effects of IDD on the sensitivity of recipients were dependent on the direction of IDD, we included the direction of IDD (negative or positive) and IDD × direction of IDD interaction in the above LMMs as explanatory variables. The identities of the interacting pairs of community members and year were specified as random intercepts. Models were generated for all three pesticide treatments. The importance of each explanatory variable was evaluated by the Type III likelihood ratio *χ*^2^ computed by the package ‘afex’ Version 0.28.0 (Singmann *et al*. 2020).

## Acknowledgements

Dr. Masayuki Ushio (HKUST) assisted greatly with our EDM analyses. We are particularly grateful to the members of the Laboratory for Conservation Ecology of Kindai University for helping with our field experiment and survey. We also thank Dr. Robin J. Smith (Lake Biwa Museum) for identifying zooplankton crustaceans, especially ostracods. The authors wish to thank Dr. Tadao Kitagawa (Kindai Univ.) for valuable technical advice. The members of the Biodiversity Assessment and Projection Section and the Climate Change Impacts Assessment Research Section of the NIES greatly helped us by joining discussions. The present study was supported by the Environment Research and Technology Development Fund (ERTDF) FY2017 (4-1701) of the Ministry of the Environment, Japan, and the Japan Society for the Promotion of Science (JSPS) KAKENHI (Grant Numbers 20K15640 and 21J01194 to K. Hashimoto and 21K18318 to D. Hayasaka and T. Kadoya).

## Supporting information

Additional Supporting Information may be found in the online version of this article.

## Data availability

Data will be available after the acceptance of this manuscript.

## Code availability

Code will be available at https://github.com/KoyaHashimoto/PaddyInteractionVariability.

## Author contributions

K.H., D.H., and T.K. conceived and designed the study. Y.E., Y.S., J.C., and K.H. collected the data. K.H. analysed the results and prepared the figures and tables. K.H., D.H., and T.K. led the writing of the manuscript. K.H., D.H., Y.E., Y.S., J.C., K.S., K.G., and T.K. contributed to revising the earlier draft. All authors have given final approval to submit this manuscript and agree to be accountable for the aspects of the work that they conducted.

## References

Abrams, P.A. (2010). Implications of flexible foraging for interspecific interactions: lessons from simple models. Funct. Ecol., 24, 7–17.

Allesina, S. & Tang, S. (2012). Stability criteria for complex ecosystems. Nature, 483, 205–208.

Barabás, G., D’Andrea, R. & Stump, S.M. (2018). Chesson’s coexistence theory. Ecol. Monogr., 88, 277–303.

Barraquand, F., Picoche, C., Detto, M. & Hartig, F. (2021). Inferring species interactions using Granger causality and convergent cross mapping. Theor. Ecol., 14, 87–105.

Bartley, T.J., McCann, K.S., Bieg, C., Cazelles, K., Granados, M., Guzzo, M.M., et al. (2019). Food web rewiring in a changing world. *Nat*. Ecol. Evol., 3, 345–354.

Baskerville, E.B. & Cobey, S. (2017). Does influenza drive absolute humidity? Proc. Natl. Acad. Sci. U. S. A., 114, E2270–E2271.

Benincá, E., Huisman, J., Heerkloss, R., Jöhnk, K.D., Branco, P., Van Nes, E.H., et al. (2008). Chaos in a long-term experiment with a plankton community. Nature, 451, 822–825.

Berlow, E.L. (1999). Strong effects of weak interactions in ecological communities. Nature, 398, 330–334.

Breheny, P. & Burchett, W. (2017). Visualization of regression models using visreg. R J., 9, 56.

Brooks, M.E., Kristensen, K., van Benthem, K.J., Magnusson, A., Berg, C.W., Nielsen, A., et al. (2017). glmmTMB balances speed and flexibility among packages for zero-inflated generalized linear mixed modeling. R J., 9, 378–400.

Calizza, E., Costantini, M.L., Careddu, G. & Rossi, L. (2017). Effect of habitat degradation on competition, carrying capacity, and species assemblage stability. Ecol. Evol., 7, 5784–5796.

Cenci, S. & Saavedra, S. (2019). Non-parametric estimation of the structural stability of non-equilibrium community dynamics. *Nat*. Ecol. Evol., 3, 912–918.

Cenci, S., Sugihara, G. & Saavedra, S. (2019). Regularized S-map for inference and forecasting with noisy ecological time series, 2019, 650–660.

Chang, C.-W., Ushio, M. & Hsieh, C. (2017). Empirical dynamic modeling for beginners. Ecol. Res., 32, 785–796.

Chang, C.W., Miki, T., Ushio, M., Ke, P.J., Lu, H.P., Shiah, F.K., et al. (2021). Reconstructing large interaction networks from empirical time series data. Ecol. Lett., 1–12.

Chesson, P. (2000). Mechanisms of maintenance of species diversity. Annu. Rev. Ecol. Syst., 31, 343–366.

Chesson, P. (2018). Updates on mechanisms of maintenance of species diversity. J. Ecol., 106, 1773–1794.

Clark, A.T., Ye, H., Isbell, F., Deyle, E.R., Cowles, J., Tilman, G.D., et al. (2015). Spatial convergent cross mapping to detect causal relationships from short time series. Ecology, 96, 1174–1181.

Cobey, S. & Baskerville, E.B. (2016). Limits to causal inference with state-space reconstruction for infectious disease. PLoS One, 11, 1–22.

Deyle, E.R., May, R.M., Munch, S.B. & Sugihara, G. (2016). Tracking and forecasting ecosystem interactions in real time. Proc. R. Soc. B Biol. Sci., 283, 20152258.

Doak, D.F., Estes, J.A., Halpern, B.S., Jacob, U., Lindberg, D.R., Lovvorn, J., et al. (2008). Understanding and predicting ecological dynamics: Are major surprises inevitable? Ecology, 89, 952–961.

Dunne, J.A. (2006). The network structure of food webs. In: Ecological Networks: Linking Structure to Dynamics in Food Webs (eds. Pascual, M. & Dunne, J.A.). Oxford University Press, New York, pp. 27–86.

Fussmann, K.E., Schwarzmüller, F., Brose, U., Jousset, A. & Rall, B.C. (2014). Ecological stability in response to warming. Nat. Clim. Chang., 4, 206–210.

Gant, D.B., Chalmers, A.E., Wolff, M.A., Hoffman, H.B. & Bushey, D.F. (1998). Fipronil: action at the GABA receptor. In: Pesticides and the Future (eds. Kuhr, R.J. & Motoyama, N.). IOS Press, Amsterdam, pp. 147–156.

Hallett, L.M., Aoyama, L., Barabás, G., Gilbert, B., Larios, L., Shackelford, N., et al. (2023). Restoration ecology through the lens of coexistence theory. Trends Ecol. Evol., 38, 1085–1096.

Hashimoto, K., Eguchi, Y., Oishi, H., Tazunoki, Y., Tokuda, M., Sánchez-Bayo, F., et al. (2019). Effects of a herbicide on paddy predatory insects depend on their microhabitat use and an insecticide application. Ecol. Appl., 29, e01945.

Hayasaka, D., Korenaga, T., Sánchez-Bayo, F. & Goka, K. (2012). Differences in ecological impacts of systemic insecticides with different physicochemical properties on biocenosis of experimental paddy fields. Ecotoxicology, 21, 191–201.

Hedges, L. V, Gurevitch, J. & Curtis, P.S. (1999). The meta-analysis of response ratio in experimental ecology. Ecology, 80, 1150–1156.

Hirai, K., Yano, T., Ugai, S., Yoshimura, T. & Hori, M. (2001). Development of a herbicide, pentoxazone. J. Pestic. Sci., 26, 194–202.

Holland, J.N., DeAngelis, D.L. & Bronstein, J.L. (2002). Population dynamics and mutualism: Functional responses of benefits and costs. Am. Nat., 159, 231–244.

Holland, J.N., Okuyama, T. & DeAngelis, D.L. (2006). Comment on “Asymmetric coevolutionary networks facilitate biodiversity maintenance”. Science, 313, 29–31.

Holling, C.S. (1959). Some characteristics of simple types of predation and parasitism. Can. Entomol., 91, 385–393.

Hsieh, C., Anderson, C. & Sugihara, G. (2008). Extending nonlinear analysis to short ecological time series. Am. Nat., 171, 71–80.

Ishiwaka, N., Hashimoto, K., Hiraiwa, M.K., Kadoya, T. & Hayasaka, D. (2024). Can warming accelerate the decline of Odonata species in experimental paddies due to insecticide fipronil exposure? Environ. Pollut., 341, 122831.

Ives, A.R., Carpenter, S.R. & Dennis, B. (1999). Community interaction webs and zooplankton responses to planktivory manipulations. Ecology, 80, 1405–1421.

Ives, A.R., Dennis, B., Cottingham, K.L. & Carpenter, S.R. (2003). Estimating community stability and ecological interactions from time-series data. Ecol. Monogr., 73, 301–330.

Japan Plant Protection Association (JPPA). (2011). Pesticide handbook 2011. Japan Plant Protection Association, Tokyo, Japan. (in Japanese)

Kadoya, T., Gellner, G. & McCann, K.S. (2018). Potential oscillators and keystone modules in food webs. Ecol. Lett., 21, 1330–1340.

Kadoya, T. & McCann, K.S. (2015). Weak interactions and instability cascades. Sci. Rep., 5, 12652.

Kasai, A., Hayashi, T.I., Ohnishi, H., Suzuki, K., Hayasaka, D. & Goka, K. (2016). Fipronil application on rice paddy fields reduces densities of common skimmer and scarlet skimmer. Sci. Rep., 6, 23055.

Kawatsu, K. (2020). Ecology and evolution of density-dependence. In: Diversity of Functional Traits and Interactions: Perspectives on Community Dynamics (ed. Mougi, A.). Springer, Singapore, pp. 161–174.

Kawatsu, K. & Kishi, S. (2018). Identifying critical interactions in complex competition dynamics between bean beetles. Oikos, 127, 553–560.

Kawatsu, K. & Kondoh, M. (2018). Density-dependent interspecific interactions and the complexity – stability relationship. Proc. R. Soc. B Biol. Sci., 285, 20180698.

Kawatsu, K., Ushio, M., van Veen, F.J.F. & Kondoh, M. (2021). Are networks of trophic interactions sufficient for understanding the dynamics of multi-trophic communities? Analysis of a tri-trophic insect food-web time-series. Ecol. Lett., 24, 543–552.

Kondoh, M. (2003). Foraging adaptation and the relationship between food-web complexity and stability. Science, 299, 1388–1391.

Kondoh, M., Kawatsu, K., Osada, Y. & Ushio, M. (2020). A data-driven approach to complex ecological systems. In: Theoretical Ecology: concepts and applications (eds. McCann, K.S. & Gellner, G.). Oxford University Press, New York, NY, USA, pp. 116–133.

Lenth, R. V. (2020). emmeans: Estimated Marginal Means, aka Least-Squares Means.

Loeuille, N. (2010). Consequences of adaptive foraging in diverse communities. Funct. Ecol., 24, 18–27.

Martinson, H.M. & Raupp, M.J. (2013). A meta-analysis of the effects of urbanization on ground beetle communities. Ecosphere, 4, 1–24.

Matsuzaki, S.S., Suzuki, K., Kadoya, T., Nakagawa, M. & Takamura, N. (2018). Bottom-up linkages between primary production, zooplankton, and fish in a shallow, hypereutrophic lake. Ecology, 99, 2025–2036.

May, R.M. (1972). Will a large complex system be stable? Nature, 238, 413–414.

McCann, K., Hastings, A. & Huxel, G.R. (1998). Weak trophic interactions and the balance of nature. Nature, 395, 794–798.

McCann, K.S. (2000). The diversity–stability debate. Nature, 405, 228–233.

McMeans, B.C., McCann, K.S., Humphries, M., Rooney, N. & Fisk, A.T. (2015). Food web structure in temporally-forced ecosystems. Trends Ecol. Evol., 30, 662–672.

Munch, S.B., Brias, A., Sugihara, G. & Rogers, T.L. (2020). Frequently asked questions about nonlinear dynamics and empirical dynamic modelling. ICES J. Mar. Sci., 77, 1463–1479.

Munch, S.B., Rogers, T.L. & Sugihara, G. (2022). Recent developments in empirical dynamic modelling. Methods Ecol. Evol., 14, 732–745.

Nagai, T., Ishihara, S., Yokoyama, A. & Iwafune, T. (2011). Effects of four rice paddy herbicides on algal cell viability and the relationship with population recovery. Environ. Toxicol. Chem., 30, 1898–1905.

Navarrete, S.A. & Berlow, E.L. (2006). Variable interaction strengths stabilize marine community pattern. Ecol. Lett., 9, 526–536.

Norris, K. & Johnstone, I. (1998). Interference competition and the functional response of oystercatchers searching for cockles by touch. Anim. Behav., 56, 639–650.

Novak, M., Yeakel, J.D., Noble, A.E., Doak, D.F., Emmerson, M., Estes, J.A., et al. (2016). Characterizing species interactions to understand press perturbations: What is the community matrix? Annu. Rev. Ecol. Evol. Syst., 47, 409–432.

O’Gorman, E.J. & Emmerson, M.C. (2009). Perturbations to trophic interactions and the stability of complex food webs. Proc. Natl. Acad. Sci. U. S. A., 106, 13393– 13398.

Oaten, A. & Murdoch, W.W. (1975). Functional response and stability in predator-prey systems. Am. Nat., 109, 289–298.

Ohgushi, T., Schmitz, O. & Holt, R.D. (Eds.). (2012). Trait-Mediated Indirect Interactions: Ecological and Evolutionary Perspectives. Cambridge University Press, Cambridge, U.K.

R Core Team. (2020). R: A language and environment for statistical computing. R Foundation for Statistical Computing.

Relyea, R.A. & Hoverman, J. (2006). Assessing the ecology in ecotoxicology: A review and synthesis in freshwater systems. Ecol. Lett., 9, 1157–1171.

Rogers, T.L., Munch, S.B., Stewart, S.D., Palkovacs, E.P., Giron-Nava, A., Matsuzaki, S. ichiro S., et al. (2020). Trophic control changes with season and nutrient loading in lakes. Ecol. Lett., 23, 1287–1297.

Rohr, J.R., Kerby, J.L. & Sih, A. (2006). Community ecology as a framework for predicting contaminant effects. Trends Ecol. Evol., 21, 606–613.

Rooney, N., McCann, K., Gellner, G. & Moore, J.C. (2006). Structural asymmetry and the stability of diverse food webs. Nature, 442, 265–269.

Rooney, N., McCann, K.S. & Moore, J.C. (2008). A landscape theory for food web architecture. Ecol. Lett., 11, 867–881.

Schielzeth, H. (2010). Simple means to improve the interpretability of regression coefficients. Methods Ecol. Evol., 1, 103–113.

Shoemaker, L.G., Sullivan, L.L., Donohue, I., Cabral, J.S., Williams, R.J., Mayfield, M.M., et al. (2020). Integrating the underlying structure of stochasticity into community ecology. Ecology, 101, e02922.

Simon-Delso, N., Amaral-Rogers, V., Belzunces, L.P., Bonmatin, J.M., Chagnon, M., Downs, C., et al. (2015). Systemic insecticides (Neonicotinoids and fipronil): Trends, uses, mode of action and metabolites. Environ. Sci. Pollut. Res., 22, 5–34.

Sugihara, G., Deyle, E.R. & Ye, H. (2017). Misconceptions about causation with synchrony and seasonal drivers. Proc. Natl. Acad. Sci. U. S. A., 114, E2272– E2274.

Sugihara, G., May, R., Ye, H., Hsieh, C.H., Deyle, E., Fogarty, M., et al. (2012). Detecting causality in complex ecosystems. Science, 338, 496–500.

Suttle, K.B., Thomsen, M.A. & Power, M.E. (2007). Species interactions reverse grassland responses to changing climate. Science, 315, 640–642.

Suzuki, K., Yoshida, K., Nakanishi, Y. & Fukuda, S. (2017). An equation-free method reveals the ecological interaction networks within complex microbial ecosystems. Methods Ecol. Evol., 8, 1774–1785.

Travis, C.C. & Post, W.M. (1979). Dynamics and comparative statics of mutualistic communities. J. Theor. Biol., 78, 553–571.

Tylianakis, J.M., Didham, R.K., Bascompte, J. & Wardle, D.A. (2008). Global change and species interactions in terrestrial ecosystems. Ecol. Lett., 11, 1351–1363.

Ushio, M., Hsieh, C., Masuda, R., Deyle, E.R., Ye, H., Chang, C.-W., et al. (2018). Fluctuating interaction network and time-varying stability of a natural fish community. Nature, 554, 360–363.

Ushio, M., Sado, T., Fukuchi, T., Sasano, S., Masuda, R., Osada, Y., et al. (2023). Temperature sensitivity of the interspecific interaction strength of coastal marine fish communities. Elife, 12, RP85795.

Uszko, W.O., Diehl, S.E., Pitsch, N.A., Lengfellner, K.A. & Muller, T. (2015). When is a type III functional response stabilizing? Theory and practice of predicting plankton dynamics under enrichment. Ecology, 96, 3243–3256.

Wang, J.Y., Kuo, T.C. & Hsieh, C. (2020). Causal effects of population dynamics and environmental changes on spatial variability of marine fishes. Nat. Commun., 11, 2635.

Wootton, J.T. & Emmerson, M. (2005). Mesurement of interaction strength in nature. Annu. Rev. Ecol. Evol. Syst., 36, 419–444.

Yamamichi, M., Gibbs, T. & Levine, J.M. (2022). Integrating eco-evolutionary dynamics and modern coexistence theory. Ecol. Lett., 25, 2091–2106.

Ye, H., Clark, A., Deyle, E. & Munch, S. (2020). rEDM: Applications of Empirical Dynamic Modeling from Time Series.

