## Supplemental materials for "Multifaceted effects of variable biotic interactions on population stability in complex interaction webs"

**Additional Supporting Information of**

**Contents**

**APPENDIX 1: Monitoring methods for paddy community members**

**APPENDIX 2: Empirical dynamic modelling**

**APPENDIX 3: Biological interpretations of the observed interaction networks**

**APPENDIX 4: Supplementary tables and figures**

Table S1. Representative taxa of the paddy community members.

Table S2. The effects of pesticide treatments on the standardized density of each community member based on linear mixed models.

Table S3. Summary of convergent cross-mapping.

Table S4. Embedding dimension and nonlinearity of each single-variable and multivariate embedding.

Fig. S1. Temporal dynamics of pesticides in the experimental mesocosms.

Fig. S2. Reconstructed interaction networks of each pesticide treatment by EDM analyses.

Fig. S3. Time series of standardized densities of paddy community members and interaction effects (represented by S-map coefficients) of all the treatments and replicates.

Fig. S4. Correlations between mean interaction strength and interaction temporal variability.

Fig. S5. Contribution of the source of variation to the values of S-map coefficients evaluated by mean squares.

**APPENDIX 1: Monitoring methods for paddy community members**

During the experiment, we collected time-series data on the densities of ten paddy community members: eukaryotic phytoplankton, rotifers, crustacean zooplankton, macrophytes, and aquatic macroinvertebrates. Aquatic macroinvertebrates were further divided into the following categories: detritivorous insects, herbivorous insects, phytophilous (clinging to macrophyte stems or leaves) predatory insects, benthic (living on bottom sediment) predatory insects, neustonic (living on the surface of water) predatory insects, and molluscs. The biological information is described in Table S1. We monitored the density of each community member every two weeks throughout the approximately 140-day experimental period until harvest in each experimental year. As such, we acquired 8 tanks × 3 years = 24 fragments of time series, each of which had 10 consecutive timepoints, for each of the 10 community members. The monitoring methods used were described previously by Hashimoto *et al.* (2019). We provide a brief description of the methods below.

To monitor the abundance of phytoplankton, rotifers, and crustacean zooplankton, we collected 500 mL water samples from 10 random sampling points in each experimental paddy. The samples were filtered through a 20-μm plankton net (Tanaka Sanjiro Co., Ltd., Fukuoka, Japan). We preserved the plankton in 5% acid Lugol's solution and then identified and counted them using an optical microscope (Olympus BX51, 100x, Olympus Corp., Tokyo, Japan) and a Sedgewick-Rafter counting chamber (Pyser-SGI Limited, Edenbridge, UK). Because most zooplankton cannot prey upon long-chained colonies of cyanobacteria, we used only eukaryotic phytoplankton data for phytoplankton counting. Note that we used different sampling protocols for crustacean zooplankton in 2017 and 2018‑2019. In 2017, we sampled water (1 L) using the same method described above but filtered the samples through a 250-μm plankton net (RIGOSHA & Co., Ltd., Tokyo, Japan). Then, the zooplankton were preserved in 4% formalin and counted using a stereoscopic microscope

(SMZ1500, Nikon Instech Co., Ltd., Tokyo, Japan). We monitored macrophyte density by setting three permanent quadrats (30 × 30 cm) in each mesocosm. We divided each quadrat into 36 grids (5 × 5 cm), counted the number of grids covered by a species and divided by the total number of grids (36). The total density of aquatic macrophytes was calculated as the sum of the coverage of each plant species. We collected aquatic macroinvertebrates by scooping a fishnet (1 mm mesh size) between the edges of each tank and rice seedlings (along a permanent transect). To reduce potential sampling errors and/or bias, fishnet scooping was conducted by the same person (Yuji Eguchi) throughout the experiment. We preserved specimens in 70% ethanol and then identified the species or closest taxonomic level and counted them.

**APPENDIX 2: Empirical dynamic modelling**

To assess the effects of pesticide application on biotic interactions within the communities in the experimental paddies, we adopted the empirical dynamic modelling (EDM) approach. The essence of EDM is a reconstruction of the true dynamics of a system by a subset of the variables of that system, which is called state-space reconstruction or attractor reconstruction. This approach is ensured by the application of Takens' embedding theorem (Takens 1981), which states that the true dynamics of a given system can be reconstructed by a set of time-lagged coordinates from a single variable of the system. Note that this theorem can be extended to cases in which there are multiple, not singular, observable variables. That is, we can reconstruct an attractor based on a subset containing several variables of a system. Collectively, by using the EDM framework, we can draw inferences about the true dynamics of a given system by observing only a subset of variables from that system, even if that system comprises many variables, such as complex biological communities.

The typical EDM procedure used to analyse interaction networks within biological communities is as follows: (1) detect dynamic causality by performing convergent cross-mapping (CCM) to determine interacting pairs of organisms and their interaction directions, and (2) perform multivariate S-map analysis to determine time-varying interaction effects among interacting pairs identified by CCM. We followed this procedure to analyse the interaction networks within communities in the experimental paddies. Below, we describe step-by-step procedures for CCM and multivariate S-map.

*Detection of causalities by CCM*

To prepare for performing the CCM and multivariate S-map analyses, we first determined the optimal embedding dimension $E$ of each community member, which is the embedding dimension that shows the best performance of the EDM predictions. The embedding dimension is the number of (time-lagged) coordinates used for state-space reconstruction. Specifically, we performed the simplex projection, which is one of the EDM methods for near-future predictions, for the whole time series of each community member, which is a composite of the data of all the tanks and every year, repeatedly by using embedding dimensions from 2 to 6. When reconstructing attractors, we concatenated fragments of time series following a previously suggested approach (Hsieh *et al.* 2008; Clark *et al.* 2015; in detail, see the paragraph below for the CCM method). Then, we compared the prediction skills of the simplex projection among different embedding dimensions. Leave-one-out cross-validation (LOOCV) was used to determine the prediction skill. The prediction skill was judged by the Pearson correlation coefficient ($\rho$), mean absolute error (MAE) and root mean squared error (RMSE). These three criteria were consistent in most cases, but if they were not consistent, we used the criterion that graphically showed the most obvious peak form.

Next, to determine if one community member had a dynamic causality on another, we performed a CCM analysis (Sugihara *et al.* 2012). CCM is a recently developed analysis used to test causalities between two variables that potentially interact with each other. As mentioned above, Takens' theorem states that attractor manifolds reconstructed by appropriate embedding preserve topological characteristics of the attractor of the whole system to which a variable used by the embedding belongs. As such, one can test whether two variables belong to the same dynamical system by comparing the topology of the reconstructed attractor manifolds of those variables. Furthermore, if variable A has a dynamic causation on variable B, a reconstructed attractor of variable B has complete information about variable A, but the reverse is not true. Thus, by examining the correspondence of the topology of the reconstructed attractors between variables A and B, we can infer not only whether there is causality between these variables but also whether the causal direction is from variables A to B, B to A, or bidirectional. Applying this principle, CCM examines the correspondence among reconstructed attractor manifolds to test causalities among variables.

We tested the causalities of all pairs of 10 community members (eukaryotic phytoplankton, rotifers, crustacean zooplankton, macrophytes, detritivorous insects, herbivorous insects, phytophilous predatory insects, benthic predatory insects, neustonic predatory insects, and molluscs). Specifically, first, we reconstructed an attractor manifold of each organism by using its best embedding dimension $E$. Although our time series had only 10 consecutive timepoints, state-space reconstruction sufficiently informative to perform CCM can be achieved by the ensemble of time series from different paddies and different years; thus, our analysis was performed with 10 timepoints × 8 paddies × 3 years = 240 timepoints in total (see Hsieh *et al.* 2008; Clark *et al.* 2015). A total of 232 timepoints were available for macrophytes because we monitored the macrophytes 9 times, from the 4th week until the 20th week (not 10 times) in 2019. When reconstructing attractors, we allowed one reconstructed attractor to consist of all the data for every replicate and year, while we constrained every single data point on the attractor to be generated only from the data of the same replicate in the same year. Second, we asked whether the reconstructed attractor manifold of a given community member (library member) could predict the value of another community member (target member). This method tested whether the former organism was causally affected by the latter. When performing predictions, we changed the amount of information of the reconstructed attractor ('library length' hereafter) from small to large by randomly sampling from the attractor, and we observed changes in the prediction performance ('cross-map skills' hereafter) with increasing library length. The minimum and maximum library lengths were $E + 1$ and $24 \times \{10 - (E - 1)\}$, respectively. The number of samples was 1,000. When there was a causal effect from the target organism to the library organism, we could observe a monotonic increase in cross-map skills from small to large library lengths (i.e., 'convergence'). We determined the cross-map skills by the Pearson correlation coefficient ($\rho$). Note that assembling multiple short time series to reconstruct an attractor assumes that different fragments of time series share common dynamic rules, which seemingly ignores potential differences in environmental conditions among experimental pesticide treatments. However, testing convergence requires a relatively large library length (because the relationships between cross-map skills and library length often show unimodal shapes, which cannot be distinguished with monotonic trends when the library length is too small), and in our case, such a large 'library length' was not achievable when we performed the analysis for each treatment separately.

We judged the causal effects of organism A on B when the CCM results met three criteria: (1) the difference in cross-map skills $\rho$ between the minimum and maximum library lengths ($\Delta\rho$) was greater than 0.1 (Ushio *et al.* 2018), (2) the cross-map skill at the maximum library length was significantly greater than that under the null hypothesis drawn from simulated surrogate data ($\alpha = 0.05$, although cases of $0.05 < P < 0.1$ were also considered in this study), and (3) the optimal time lag determined by lagged CCM (Ye *et al.* 2015) was not greater than 0. We tested these criteria sequentially in the order in which they are listed and proceeded to the next criterion only if the prior criterion was met.

There are several ways to simulate surrogate time series. Among these methods, we chose the 'seasonal surrogate' method, which is designed for time series with a constant cycle, such as seasons (Deyle *et al.* 2016a). In short, this method detects and preserves a cyclic trend from an observed time series while randomly shuffling anomalies from the average of this cyclic trend. This gave a 'seasonal surrogate' time series that had the same cyclic trend relative to the original time series but with random anomalies. To simulate surrogate data, first, we treated 24 fragments of time series as one consecutive time series with a 10-timepoint cycle. Second, we generated surrogate data by using the function 'surrogate_seasonal' in the package 'rEDM' Version 0.7.5 (Ye *et al.* 2020). We simulated 1,000 surrogate data points for each of the 10 organisms. Then, we drew the distribution of cross-map skills under the null hypothesis (i.e., there was no causality) and calculated the *P* values of the observed cross-map skills.

To determine the optimal time lag for cross-mapping, we performed lagged CCM (Ye *et al.* 2015). Lagged CCM alters the time lag between the library and target variables when performing predictions, and one can compare cross-map skills among different time lags. Interestingly, lagged CCM can be applied to measure cross-map skills on the future state of target variables (i.e., time lags are positive). The better cross-map skills of a positive time lag than those of the current or past states of target variables can be regarded as a signal of false positives. In the lagged CCM analysis, we simulated 100 random samples for each time lag from –2 to +2 and compared the mean values of the cross-map skills $\rho$ among the time lags.

*Tracking interaction effects by multivariate S-map*

To estimate the interaction effects among community members in the experimental paddies at each time point, we performed multivariate S-map analysis. The multivariate S-map is a locally weighted sequential linear regression at each location of an attractor manifold reconstructed by multivariate embedding. Deyle *et al.* (2016b) argued that the regression coefficients of S-map ('S-map coefficients') at each time point can be interpreted as time-varying interaction effects. That is, S-map coefficients are estimated at each time point of the time series data sequentially, yielding different values of interaction strength among different time points (i.e., 'time-varying'). This feature of S-map coefficients can be used to explore the context dependency of interaction effects (Ushio *et al.* 2018; Liu & Gaines 2022). Note that the interpretation of the S-map coefficient depends on the choice of the response variable of sequential regressions. We chose the *per capita* population growth rate as the response variable to determine the interaction strength at the *per capita* level (Suzuki *et al.* 2017). The *per capita* growth rate was approximated by calculating $\ln(N_{t+1}/N_t)$, where $N_t$ and $N_{t+1}$ are the raw abundance data of the focal species at time points $t$ and $t$+1, respectively, assuming exponential growth from time point $t$ to $t$+1. Since there were several zero values in the raw abundance data, one was added to every abundance data point to calculate the *per capita* growth rate.

First, we chose multivariate coordinates to reconstruct a multivariate attractor manifold for each recipient (i.e., causally influenced by donors) by using the community members, the causalities of which were detected by CCM. Note that the values of the S-map coefficients were sensitive to the choice of embedding elements. There are several ways to reconstruct a multivariate attractor manifold. We adopted the approach in which the elements of univariate lagged embedding of a recipient were substituted by the donor densities to ensure that the dimension of multivariate embedding was no less than $E$. For example, when the $E$ of a recipient was 5 and when three donors were detected by CCM, multivariate embedding was performed by taking the coordinates $\{N_{Rt}, N_{Rt-1}, N_{D1t}, N_{D2t}, N_{D3t}\}$, where $N_{Rt}$ is the density of the recipient at time $t$ and $N_{Dit}$ is the density of donor $i$ ($i$ = 1, 2, 3). We used the ensemble of time series from different paddies and different years to reconstruct multivariate attractors (the same as for simplex projection and CCM).

Second, we determined the optimal values of the two parameters ($\theta$ and $\lambda$) in S-map models by repeatedly performing multivariate S-map for each recipient. The parameter $\theta$ corresponds to the amount of weighting for locally weighted sequential regressions in the S-map, determining how locally the information from the whole state space is to be used for each regression. Thus, the nonlinearity of the reconstructed dynamics can be measured by $\theta$. When $\theta = 0$, the S-map becomes a nonweighted linear regression. The parameter $\lambda$ controls the L2 penalization to avoid overfitting. Cenci *et al.* (2019) recently proposed that incorporating such penalization into S-map can improve prediction skill even when using natural, ecological data with large amounts of noise (known as 'regularized S-map'), and is thus suitable for our study. We repeatedly performed a multivariate S-map to predict the *per capita* growth of recipient

$\ln(N_{Rt+1}/N_{Rt})$ by changing the $\theta$ value by (0, 0.1, 0.5, 1, 1.5, 2, 2.5, 3, 4, 6, 8) and the $\lambda$ value by (0, 0.0001, 0.001, 0.01, 0.1, 0.5, 1, 2) for each multivariate embedding. Then, we compared the prediction skills (RMSEs) among different values of $\theta$ and $\lambda$ determined by LOOCV and used the values showing the minimum value of the RMSE for further analyses.

Third, using the optimal $\theta$ and $\lambda$ values determined in the previous step, we again performed a multivariate S-map for each multivariate embedding. For embedding to predict community member B, the estimated S-map coefficients of community member A were regarded as the time-varying interaction effect of community member A on B. Note that S-map analyses were performed for reconstructed manifolds embedded by using all the data for eight paddies and three years as a composite dataset. Thus, we obtained the interaction effect not only for every time but also for every paddy (and thus every treatment) and every year.

### APPENDIX 3: Biological interpretations of the observed interaction networks

*Cascading positive effects from phytoplankton to crustacean zooplankton*

Phytoplankton may be food resources for rotifers, and rotifers may be foraged by larger crustacean zooplankton. Negative effects of higher trophic levels were observed only for rotifers to phytoplankton, suggesting that bottom-up effects were more prevalent in these mesocosms. Indeed, paddy systems are more likely than other freshwater systems to be donor-controlled (Okuda 2012).

*Phytoplankton had negative effects on detritivores and herbivores*

Phytoplankton may negatively affect freshwater biodiversity by modifying water conditions, leading to conditions such as eutrophication and increased turbidity (Scheffer *et al.* 2001). The observed negative effects on these macroinvertebrates may reflect such negative effects on water conditions.

*Macrophytes had a positive effect on neustonic predators*

Macrophytes gather small arthropods, such as aphids and other hemipterans, above the water surface. These small arthropods occasionally drop onto the water surface and may serve as food resources for neustonic predators (water striders).

*Bidirectional effects between rotifers and phytophilous predators*

Bidirectional effects between rotifers and phytophilous predators strongly suggest a prey–predator interaction. Phytophilous predators, i.e., damselfly and dragonfly larvae, are known to prey upon rotifers, especially in their earlier stages (Sugita *et al.* 2018). Thus, the negative effects of phytophilous predators on rotifers are reasonable. In addition, the positive effects of phytophilous predators on phytoplankton suggest top-down trophic cascades via the predation of rotifers by phytophilous predators. On the other hand, because phytophilous predators were able to escape from the mesocosms after their development was complete, the observed negative effects of rotifers on phytophilous predators (in I, H, and I+H) do not necessarily mean that rotifers have negative effects on the survival, development, and fitness of phytophilous predators.

*Negative effects of benthic predators on molluscs*

Benthic predators (dragonfly larvae) may prey upon young molluscs (snails and clams). This may be reflected by the apparent positive effects of the insecticide (fipronil) on mollusc density because benthic predators were strongly negatively affected by insecticide application, suggesting density-mediated indirect effects of the insecticide on molluscs by decreasing predation by benthic predators on molluscs.

### APPENDIX 4: Supplementary tables and figures

Table S1. Representative taxa of the paddy community members.

| Community members | Representative taxa |
|---|---|
| Eukaryotic phytoplankton | *Cryptomonas* spp. (algae) |
| | *Euglena* spp. (phyotosynthetic flagellates) |
| | *Cosmarium* spp. (algae) |
| | Pennate diatoms (diatoms) |
| Rotifers | Brachionidae |
| | Lecanidae |
| | Synchaetidae |
| Crustacean zooplankton | Chydoridae (cradocerans) |
| | Ostracoda |
| | Copepoda |
| Macrophytes | *Monochoria vaginalis* (pondweed) |
| | *Schoenoplectiella hotarui* (sedge) |
| | *Azolla* sp. (water fern) |
| Detritivorous insects | Chironomidae (nonbiting midges) |
| | *Aedes* spp. (mosquitos) |
| | *Anopheles* spp. (mosquitos) |
| Herbivorous insects | Corixidae spp. (water boatmen) |
| | Curculionoidea spp. (weevils) |
| | Pyraloidea spp. (snout moths) |
| Phytophilous predatory insects | *Indolestes peregrinus* (damselfly) |
| | *Cercion calamorum* (damselfly) |
| | *Anax parthenope* (dragonfly) |
| Benthic predatory insects | *Crocothemis servilia* (dragonfly) |
| | *Orthetrum albistylum* (dragonfly) |
| Neustonic predatory insects | *Microvelia douglasi* (small water strider) |
| | *Gerris gracilicornis* (water strider) |
| Molluscs | *Physa acuta* (snail) |
| | Bivalvia spp. (freshwater clams) |

Table S2. The effects of pesticide treatments on the standardized density of each community member based on linear mixed models. For fixed effects (treatment, week, and treatment × week interaction), the results of Type III likelihood ratio tests are shown. For random effects (AR1 temporal autocorrelation, tank identity, year, and residuals), the estimated autocorrelation coefficient $\rho$ or variance $\sigma^2$ is shown. Bold indicates statistical significance.

| | Treatment | | | Week | | | Treatment × Week | | |
|---|---|---|---|---|---|---|---|---|---|
| | LR-$\chi^2$ | df | *P* | LR-$\chi^2$ | df | *P* | LR-$\chi^2$ | df | *P* |
| Eukaryotic phytoplankton | 8.24 | 3 | **0.04** | 34.64 | 9 | **< 0.001** | 36.97 | 27 | 0.10 |
| Rotifers | 1.86 | 3 | 0.6 | 24.54 | 9 | **< 0.01** | 36.94 | 27 | 0.10 |
| Crustacean zooplankton | 2.89 | 3 | 0.4 | 76.86 | 9 | **< 0.001** | 29.81 | 27 | 0.3 |
| Macrophytes | 14.39 | 3 | **< 0.01** | 172.70 | 9 | **< 0.001** | 31.37 | 27 | 0.3 |
| Detritivorous insects | 5.93 | 3 | 0.1 | 44.58 | 9 | **< 0.001** | 23.22 | 27 | 0.7 |
| Herbivorous insects | 13.03 | 3 | **< 0.01** | 23.84 | 9 | **< 0.01** | 65.23 | 27 | **< 0.001** |
| Phytophilous predatory insects | 14.54 | 3 | **< 0.01** | 23.73 | 9 | **< 0.01** | 27.66 | 27 | 0.4 |
| Benthic predatory insects | 22.43 | 3 | **< 0.001** | 24.68 | 9 | **< 0.01** | 35.92 | 27 | 0.1 |
| Neustonic predatory insects | 4.86 | 3 | 0.2 | 46.82 | 9 | **< 0.001** | 30.17 | 27 | 0.3 |
| Molluscs | 11.49 | 3 | **< 0.01** | 11.62 | 9 | 0.2 | 26.78 | 27 | 0.5 |

| | AR1 $\rho$ | $\sigma^2$_Tank | $\sigma^2$_Year | $\sigma^2$_residual |
|---|---|---|---|---|
| Eukaryotic phytoplankton | 0.58 | $8.0 \times 10^{-10}$ | $2.1 \times 10^{-2}$ | 0.64 |
| Rotifers | 0.30 | 0.021 | 0.40 | 0.65 |
| Crustacean zooplankton | 0.47 | $5.1 \times 10^{-6}$ | $2.2 \times 10^{-6}$ | 0.77 |
| Macrophytes | 0.90 | $2.0 \times 10^{-4}$ | 0.10 | 0.44 |
| Detritivorous insects | 0.27 | 0.059 | 0.029 | 0.80 |
| Herbivorous insects | 0.16 | $1.3 \times 10^{-5}$ | $5.0 \times 10^{-2}$ | 0.74 |
| Phytophilous predatory insects | 0.43 | $4.9 \times 10^{-8}$ | 0.11 | 0.71 |
| Benthic predatory insects | 0.21 | 0.038 | $4.7 \times 10^{-2}$ | 0.46 |
| Neustonic predatory insects | 0.40 | 0.075 | 0.017 | 0.85 |
| Molluscs | 0.42 | 0.055 | 0.51 | 0.48 |

Table S3. Summary of convergent cross-mapping. The rows and columns represent recipient and donor community members, respectively. $\Delta\rho$: the differences in cross-map skills ($\rho$) at the minimum and the maximum library lengths. $P$: estimated $P$ values based on the surrogate time series analysis. Optimal lag: the time lag between the time series of donor and recipient organisms that showed the best cross-map skills. The bold numbers indicate the causes that met the above three criteria and thus were considered in this study.

| | Donor organisms | | | | | | | | | |
|---|---|---|---|---|---|---|---|---|---|---|
| Recipient organisms | Ph | Ro | Zo | Ma | De | He | Pp | Bp | Np | Mo |
| Phytoplankton (Ph) | - | **$\Delta\rho$ = 0.28**<br>**$P$ < 0.001**<br>**Optimal lag = –1** | $\Delta\rho$ = 0.06 | $\Delta\rho$ = –0.12 | **$\Delta\rho$ = 0.16**<br>**$P$ = 0.07**<br>**Optimal lag = –1** | $\Delta\rho$ = –0.04 | **$\Delta\rho$ = 0.13**<br>**$P$ < 0.05**<br>**Optimal lag = –2** | $\Delta\rho$ = 0.13<br>$P$ < 0.05<br>Optimal lag = 2 | $\Delta\rho$ = 0.01 | $\Delta\rho$ = 0.07 |
| Rotifers (Ro) | **$\Delta\rho$ = 0.19**<br>**$P$ < 0.01**<br>**Optimal lag = –2** | - | $\Delta\rho$ = 0.22<br>$P$ < 0.05<br>Optimal lag = 2 | $\Delta\rho$ = –0.15 | $\Delta\rho$ = 0.006 | **$\Delta\rho$ = 0.17**<br>**$P$ < 0.01**<br>**Optimal lag = 0** | **$\Delta\rho$ = 0.23**<br>**$P$ < 0.01**<br>**Optimal lag = –2** | $\Delta\rho$ = 0.22<br>$P$ < 0.01<br>Optimal lag = 2 | $\Delta\rho$ = 0.002 | $\Delta\rho$ = 0.24<br>$P$ < 0.01<br>Optimal lag =2 |
| Crustacean zooplankton (Zo) | $\Delta\rho$ = –0.11 | **$\Delta\rho$ = 0.29**<br>**$P$ < 0.01**<br>**Optimal lag = –2** | - | $\Delta\rho$ = 0.17<br>$P$ = 0.17 | $\Delta\rho$ = 0.08 | $\Delta\rho$ = 0.25<br>$P$ < 0.01<br>Optimal lag = 2 | $\Delta\rho$ = 0.22<br>$P$ < 0.05<br>Optimal lag = 2 | $\Delta\rho$ = 0.19<br>$P$ = 0.06<br>Optimal lag = 2 | $\Delta\rho$ = 0.24<br>$P$ < 0.05<br>Optimal lag = 2 | $\Delta\rho$ = –0.05 |
| Macrophytes (Ma) | $\Delta\rho$ = 0.35<br>$P$ < 0.001<br>Optimal lag = 1 | **$\Delta\rho$ = 0.28**<br>**$P$ < 0.01**<br>**Optimal lag = –2** | $\Delta\rho$ = 0.25<br>$P$ < 0.05<br>Optimal lag = 2 | - | $\Delta\rho$ = 0.21<br>$P$ < 0.001<br>Optimal lag = 2 | **$\Delta\rho$ = 0.27**<br>**$P$ < 0.001**<br>**Optimal lag = –1** | **$\Delta\rho$ = 0.15**<br>**$P$ < 0.05**<br>**Optimal lag = –1** | $\Delta\rho$ = 0.31<br>$P$ < 0.001<br>Optimal lag = 2 | $\Delta\rho$ = 0.21<br>$P$ < 0.001<br>Optimal lag = 2 | $\Delta\rho$ = 0.12<br>$P$ < 0.05<br>Optimal lag = 1 |
| Detritivores (De) | **$\Delta\rho$ = 0.22**<br>**$P$ < 0.01**<br>**Optimal lag = 0** | **$\Delta\rho$ = 0.10**<br>**$P$ = 0.09**<br>**Optimal lag = –2** | $\Delta\rho$ = 0.18<br>$P$ = 0.18 | **$\Delta\rho$ = 0.14**<br>**$P$ < 0.01**<br>**Optimal lag = –2** | - | $\Delta\rho$ = 0.13<br>$P$ = 0.06<br>Optimal lag = 2 | $\Delta\rho$ = –0.12 | $\Delta\rho$ = 0.08 | $\Delta\rho$ = 0.07 | $\Delta\rho$ = 0.01 |
| Herbivores (He) | **$\Delta\rho$ = 0.14**<br>**$P$ < 0.05**<br>**Optimal lag = 0** | $\Delta\rho$ = 0.03 | $\Delta\rho$ = 0.08 | **$\Delta\rho$ = 0.20**<br>**$P$ < 0.01**<br>**Optimal lag = –2** | **$\Delta\rho$ = 0.21**<br>**$P$ < 0.01**<br>**Optimal lag = –2** | - | $\Delta\rho$ = 0.16<br>$P$ < 0.01<br>Optimal lag = 2 | $\Delta\rho$ = 0.33<br>$P$ < 0.001<br>Optimal lag = 2 | $\Delta\rho$ = 0.04 | $\Delta\rho$ = 0.16<br>$P$ = 0.13 |
| Phytophilous predators (Pp) | $\Delta\rho$ = –0.01 | **$\Delta\rho$ = 0.21**<br>**$P$ < 0.05**<br>**Optimal lag = –2** | $\Delta\rho$ = 0.17<br>$P$ = 0.18 | $\Delta\rho$ = –0.08 | $\Delta\rho$ = 0.03 | $\Delta\rho$ = 0.01 | - | $\Delta\rho$ = 0.05 | $\Delta\rho$ = 0.03 | $\Delta\rho$ = 0.02 |
| Benthic predators (Bp) | $\Delta\rho$ = 0.01 | $\Delta\rho$ = 0.03 | $\Delta\rho$ = 0.01 | $\Delta\rho$ = 0.003 | $\Delta\rho$ = –0.02 | $\Delta\rho$ = 0.12<br>$P$ = 0.13 | $\Delta\rho$ = –0.003 | - | $\Delta\rho$ = –0.13 | $\Delta\rho$ = 0.05 |
| Neustonic predators (Np) | $\Delta\rho$ = 0.095 | $\Delta\rho$ = 0.12<br>$P$ = 0.10 | $\Delta\rho$ = 0.11<br>$P$ = 0.30 | **$\Delta\rho$ = 0.16**<br>**$P$ < 0.001**<br>**Optimal lag = –2** | $\Delta\rho$ = 0.03 | $\Delta\rho$ = 0.06 | $\Delta\rho$ = 0.06 | $\Delta\rho$ = 0.14<br>$P$ = 0.09<br>Optimal lag = 1 | - | $\Delta\rho$ = –0.04 |
| Molluscs (Mo) | $\Delta\rho$ = 0.11<br>$P$ < 0.001<br>Optimal lag = 1 | **$\Delta\rho$ = 0.18**<br>**$P$ < 0.001**<br>**Optimal lag = –1** | **$\Delta\rho$ = 0.11**<br>**$P$ = 0.09**<br>**Optimal lag = –2** | $\Delta\rho$ = 0.10<br>$P$ = 0.11 | $\Delta\rho$ = 0.11<br>$P$ = 0.19 | **$\Delta\rho$ = 0.13**<br>**$P$ < 0.05**<br>**Optimal lag = 0** | $\Delta\rho$ = 0.17<br>$P$ < 0.05<br>Optimal lag = 1 | **$\Delta\rho$ = 0.17**<br>**$P$ < 0.001**<br>**Optimal lag = –2** | **$\Delta\rho$ = 0.27**<br>**$P$ < 0.01**<br>**Optimal lag = –1** | - |

Table S4. Embedding dimension and nonlinearity of each single-variable and multivariate embedding. *E*: Optimal embedding dimension determined by simplex projection (for multivariate embedding, the dimension actually used was reported). $\theta$: best nonlinearity parameters determined by the S-map method. $\lambda$: the L2 penalization parameter to avoid overfitting. Recipients and donors: receivers and initiators of interaction effects determined by convergent cross-mapping, respectively.

Embedding

| Single variate embedding | *E* | $\theta$ |
|---|---|---|
| Eukaryotic phytoplankton | 5 | 0.75 |
| Rotifers | 6 | 1.5 |
| Crustacean zooplankton | 6 | 1 |
| Macrophytes | 5 | 0.3 |
| Detritivorous insects | 4 | 2 |
| Herbivorous insects | 6 | 1 |
| Phytophilous predatory insects | 5 | 0.75 |
| Benthic predatory insects | 2 | 3 |
| Neustonic predatory insects | 5 | 0.03 |
| Molluscs | 4 | 0.75 |

Multivariate embedding

| Recipients | Donors | *E* | $\theta$ | $\lambda$ |
|---|---|---|---|---|
| Eukaryotic phytoplankton | Rotifers, Detritivorous insects, Phytophilous predatory insects | 5 | 2 | 0.01 |
| Rotifers | Eukaryotic phytoplankton, Herbivorous insects, Phytophilous predatory insects | 6 | 1 | 0.1 |
| Crustacean zooplankton | Rotifers | 6 | 0 | 0.1 |
| Macrophytes | Rotifers, Herbivorous insects, Phytophilous predatory insects | 5 | 6 | 0.1 |
| Detritivorous insects | Eukaryotic phytoplankton, Rotifers, Macrophytes | 4 | 2.5 | 0 |
| Herbivorous insects | Eukaryotic phytoplankton, Macrophytes, Detritivorous insects | 6 | 0 | 0.01 |
| Phytophilous predatory insects | Rotifers | 5 | 8 | 2 |
| Neustonic predatory insects | Macrophytes | 5 | 0 | 0.01 |
| Molluscs | Rotifers, Crustacean zooplankton, Herbivorous insects, Benthic predatory insects, Neustonic predatory insects | 5 | 6 | 1 |

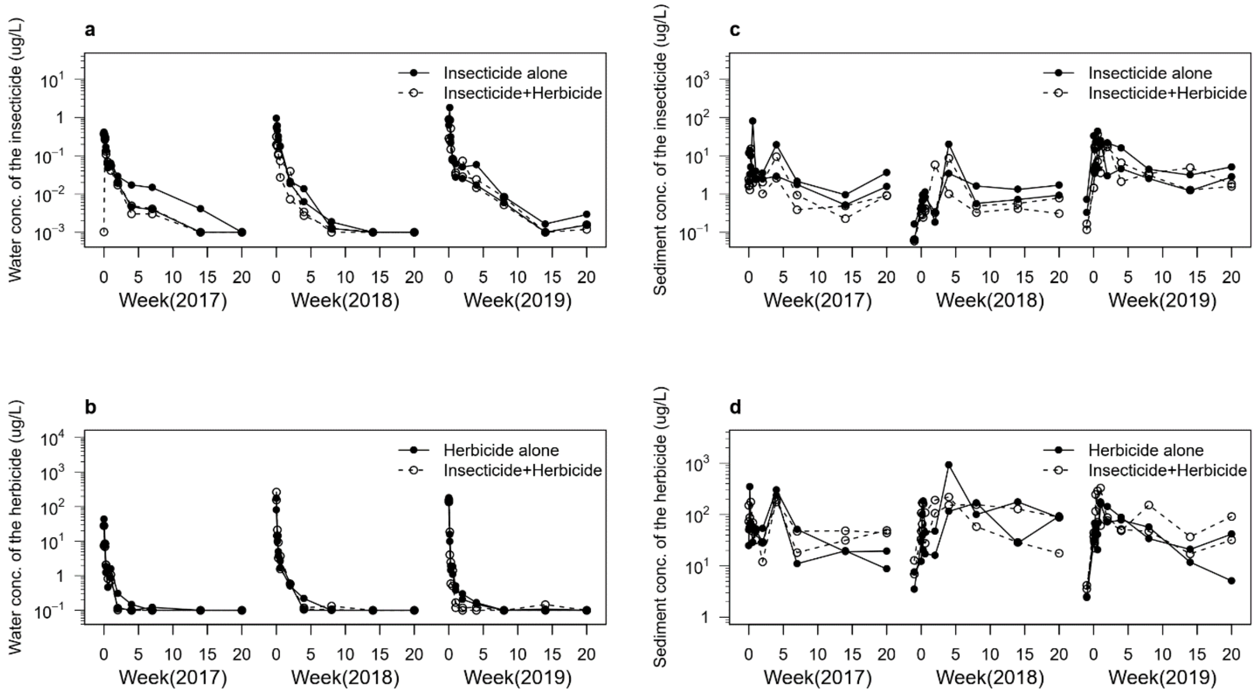


Fig. S1. Temporal dynamics of pesticides in the experimental mesocosms. **a**, **b**, Concentration of **a**) the insecticide (fipronil) and **b**) the herbicide (pentoxazone) in water. **c**, **d**, Concentration of **c**) the insecticide and **d**) the herbicide in sediment. The data below the limits of detection were substituted with limit values to facilitate visual interpretation.

Note: We collected water (50 mL) from 20 random sampling locations (2.5 mL per spot) in each pesticide-treated mesocosm. We also collected surface sediment (100 g, 2–3 cm depth) from 10 random sampling locations (10 g per spot) in each mesocosm. To avoid photolysis and degradation of both pesticides, we placed the collected samples in amber bottles sealed with aluminium foil and stored them in a refrigerator (5 °C) until analysis. Analyses of pentoxazone and fipronil in water and sediment were carried out at the certified analytical laboratories of HEISEIRIKEN Co., Ltd. (Utsunomiya, Tochigi Prefecture, Japan) using LC–MS/MS analysis.

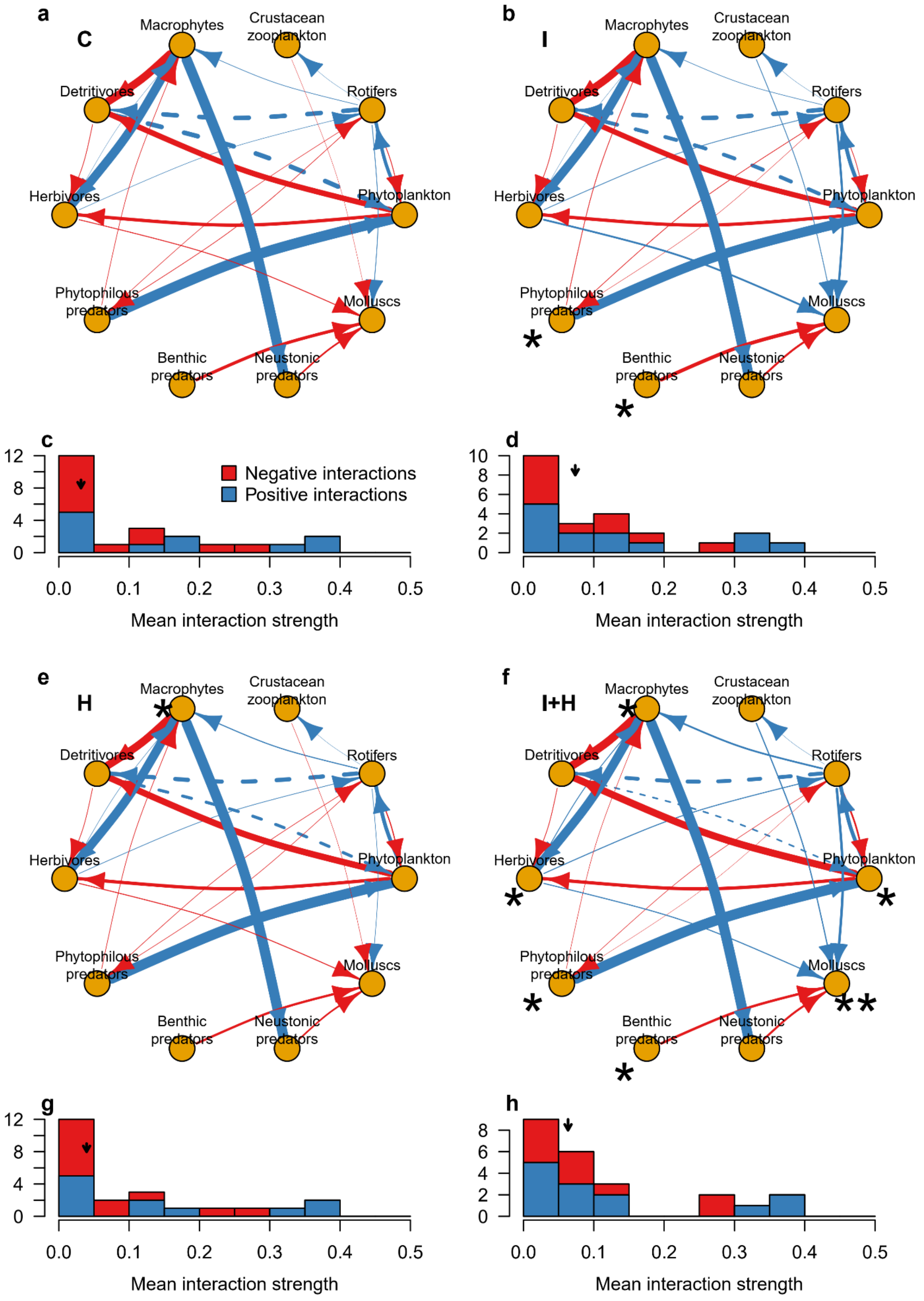
a
C
b
I
Macrophytes
Crustacean zooplankton
Detritivores
Rotifers
Herbivores
Phytoplankton
Phytophilous predators
Molluscs
Benthic predators
Neustonic predators
c
d
Negative interactions
Positive interactions
Mean interaction strength
e
H
f
I+H
g
h

Fig. S2. Reconstructed interaction networks of each pesticide treatment by EDM analyses. C: control, I: insecticide (fipronil) treatment, H: herbicide (pentoxazone) treatment, and I+H: insecticide + herbicide mixture treatment. In **a**, **b**, **e**, and **f**, the red and blue arrows indicate negative and positive interactions, respectively. Their thickness is proportional to the *per capita* interaction effects represented by the absolute values of the S-map coefficient averaged over all the experimental periods and replicates. Solid arrows: $P < 0.05$ and dashed arrows: $0.05 < P < 0.1$. One asterisk in **b**, **e** and **f** indicates that the indicated member was significantly decreased by the pesticide treatment relative to the controls, whereas two asterisks in **f** indicate that the indicated member was significantly increased by the pesticide treatment relative to the controls. In **c**, **d**, **g**, and **h**, the distributions of the absolute values of the S-map coefficient averaged over all the experimental periods and replicates are shown. The vertical arrows indicate the median values.

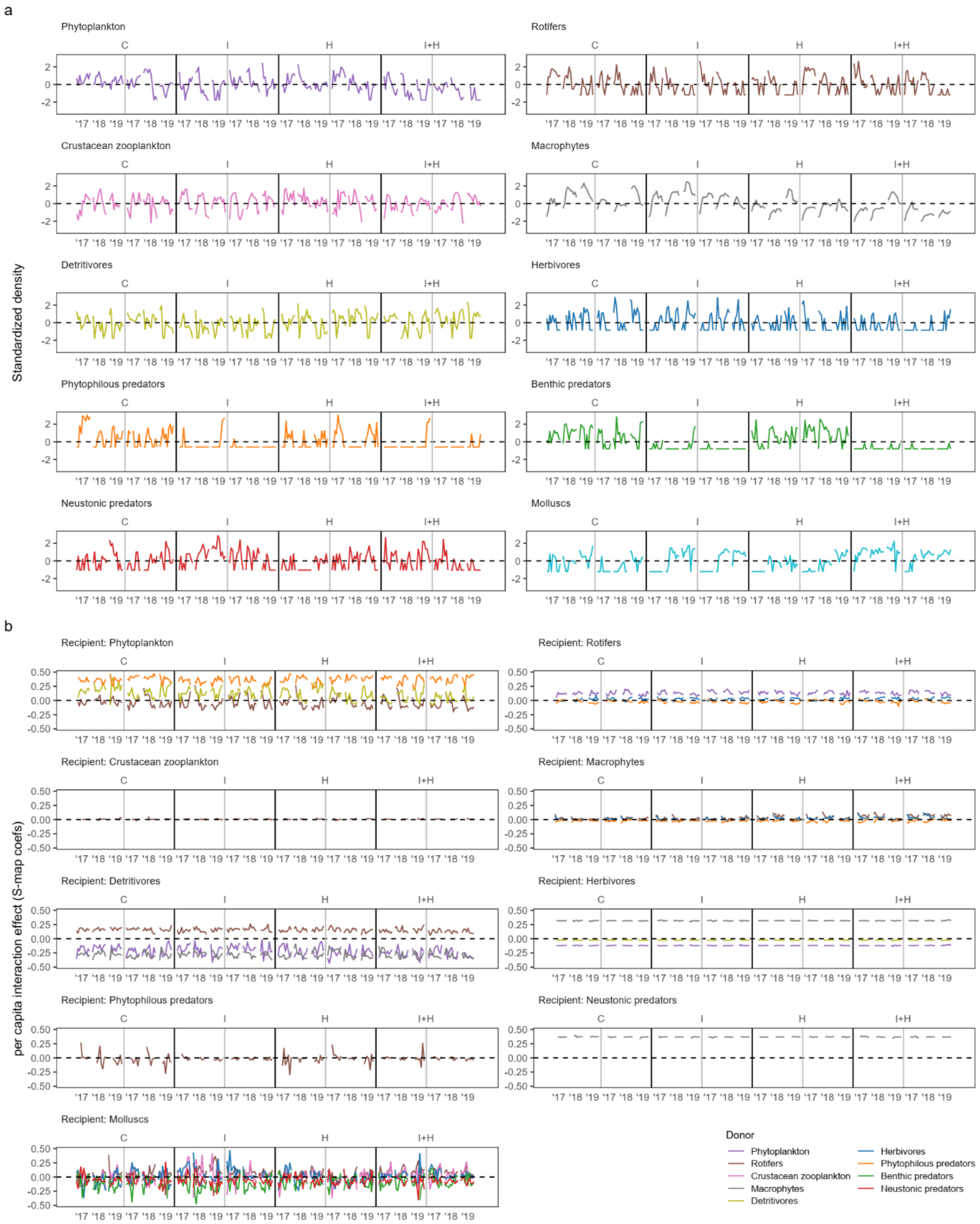


Fig. S3. Time series of **a**) standardized densities of community members in experimental paddies and **b**) *per capita* interaction effect (S-map coefficients) of all the treatments and replicates. The panels are split by interaction recipients.

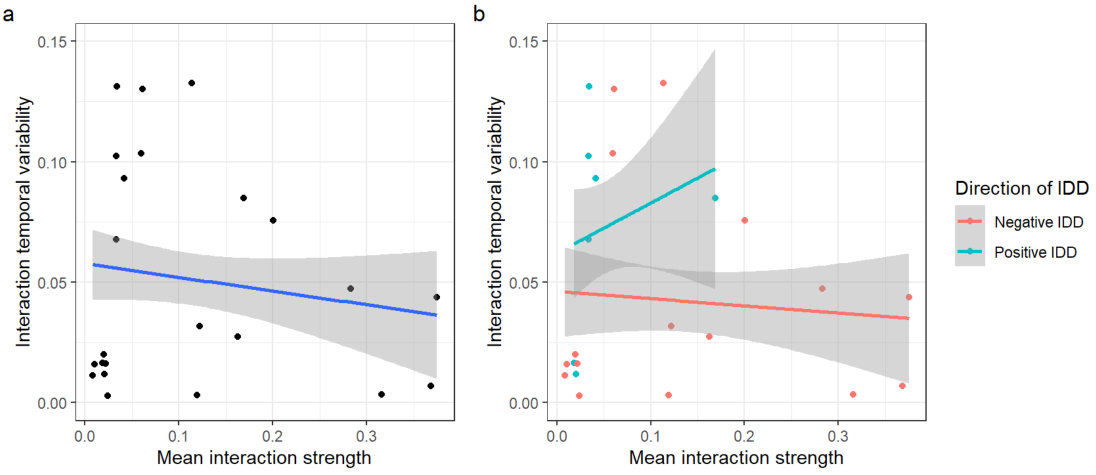


Fig. S4. Correlations between mean interaction strength and interaction temporal variability. (a) and (b) use the same data, but the latter panel explicitly shows the direction of recipient density-dependence in *per capita* interaction effect (interaction density-dependence; IDD). Although the correlation was not statistically significant (*P* = 0.2), some of the weaker interactions were temporally variable, whereas the stronger interactions were relatively temporally stable. This pattern was mainly driven by interactions with negative IDD.

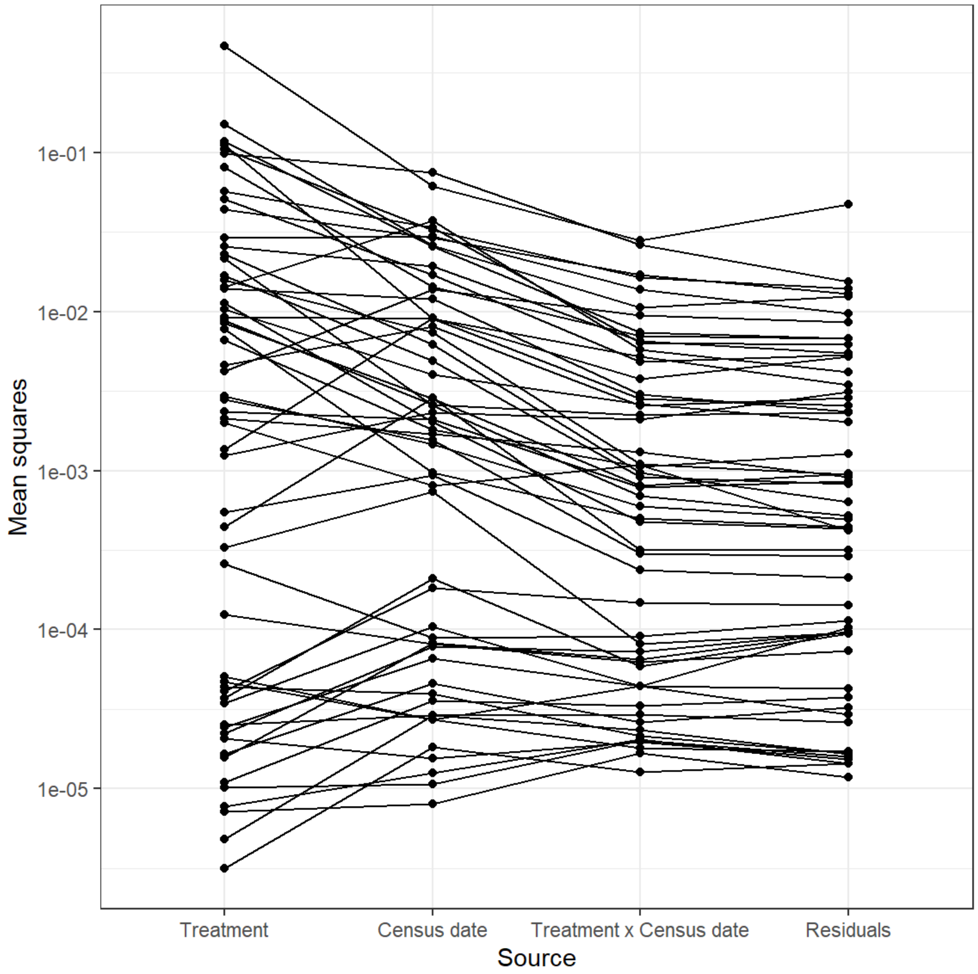


Fig. S5. Contribution of the source of variation to the values of S-map coefficients evaluated by mean squares. Each solid line represents one ANOVA model examining the effects of treatment, census week, their interaction and residuals on the value of an S-map coefficient. Of the 57 S-map coefficients, for 41 coefficients, the contribution of treatment was greater than that of residuals, and for 48 coefficients, the contribution of census date was greater than that of residuals.

### References


Cenci, S., Sugihara, G. & Saavedra, S. (2019). Regularized S-map for inference and forecasting with noisy ecological time series, 2019, 650–660.

Clark, A.T., Ye, H., Isbell, F., Deyle, E.R., Cowles, J., Tilman, G.D., *et al.* (2015). Spatial convergent cross mapping to detect causal relationships from short time series. *Ecology*, 96, 1174–1181.

Deyle, E.R., Maher, M.C., Hernandez, R.D., Basu, S. & Sugihara, G. (2016a). Global environmental drivers of influenza. *Proc. Natl. Acad. Sci. U. S. A.*, 113, 13081–13086.

Deyle, E.R., May, R.M., Munch, S.B. & Sugihara, G. (2016b). Tracking and forecasting ecosystem interactions in real time. *Proc. R. Soc. B Biol. Sci.*, 283, 20152258.

Hashimoto, K., Eguchi, Y., Oishi, H., Tazunoki, Y., Tokuda, M., Sánchez-Bayo, F., *et al.* (2019). Effects of a herbicide on paddy predatory insects depend on their microhabitat use and an insecticide application. *Ecol. Appl.*, 29, e01945.

Hsieh, C., Anderson, C. & Sugihara, G. (2008). Extending nonlinear analysis to short ecological time series. *Am. Nat.*, 171, 71–80.

Liu, O.R. & Gaines, S.D. (2022). Environmental context dependency in species interactions. *Proc. Natl. Acad. Sci. U. S. A.*, 119, 1–11.

Okuda, N. (2012). How to assess ecosystem structure and functioning of paddy fields using stable isotopes. *Japanese J. Ecol.*, 62, 207–216. (in Japanese)

Scheffer, M., Carpenter, S., Foley, J.A., Folke, C. & Walker, B. (2001). Catastrophic shifts in ecosystems. *Nature*, 413, 591–596.

Sugihara, G., May, R., Ye, H., Hsieh, C.H., Deyle, E., Fogarty, M., *et al.* (2012). Detecting causality in complex ecosystems. *Science.*, 338, 496–500.

Sugita, N., Agemori, H. & Goka, K. (2018). Acute toxicity of neonicotinoids and some insecticides to first instar nymphs of a non-target damselfly, *Ischnura senegalensis* (Odonata: Coenagrionidae), in Japanese paddy fields. *Appl. Entomol. Zool.*, 53,

519–524.

Suzuki, K., Yoshida, K., Nakanishi, Y. & Fukuda, S. (2017). An equation-free method reveals the ecological interaction networks within complex microbial ecosystems. *Methods Ecol. Evol.*, 8, 1774–1785.

Takens, F. (1981). Detecting strange attractors in turbulence. In: *Dynamical Systems and Turbulence, Warwick 1980* (eds. Rand, D. & Young, L.-S.). Springer, Berlin, Germany, pp. 366–381.

Ushio, M., Hsieh, C., Masuda, R., Deyle, E.R., Ye, H., Chang, C.-W., *et al.* (2018). Fluctuating interaction network and time-varying stability of a natural fish community. *Nature*, 554, 360–363.

Ye, H., Clark, A., Deyle, E. & Munch, S. (2020). rEDM: Applications of Empirical Dynamic Modeling from Time Series.

Ye, H., Deyle, E.R., Gilarranz, L.J. & Sugihara, G. (2015). Distinguishing time-delayed causal interactions using convergent cross mapping. *Sci. Rep.*, 5, 1–9.
